# REPROGRAMMING CBX8-PRC1 FUNCTION WITH A POSITIVE ALLOSTERIC MODULATOR

**DOI:** 10.1101/2021.02.23.432388

**Authors:** Junghyun L. Suh, Daniel Bsteh, Yibo Si, Bryce Hart, Tyler M. Weaver, Carina Pribitzer, Roy Lau, Shivani Soni, Heather Ogana, Justin M. Rectenwald, Jacqueline L. Norris, Stephanie H. Cholensky, Cari Sagum, Jessica D. Umana, Dongxu Li, Brian Hardy, Mark T. Bedford, Shannon M. Mumenthaler, Heinz-Josef Lenz, Yong-mi Kim, Gang Greg Wang, Ken H. Pearce, Lindsey I. James, Dmitri B. Kireev, Catherine A. Musselman, Stephen V. Frye, Oliver Bell

**Author notes:** Correspondence (S.V.F.), (O.B.).

## Abstract

Canonical targeting of Polycomb Repressive Complex 1 (PRC1) to repress developmental genes is mediated by cell type-specific, paralogous chromobox (CBX) proteins (CBX2, 4, 6, 7 and 8). Based on their central role in silencing and their misregulation associated with human disease including cancer, CBX proteins are attractive targets for small molecule chemical probe development. Here, we have used a quantitative and target-specific cellular assay to discover a potent positive allosteric modulator (PAM) of CBX8. The PAM activity of UNC7040 antagonizes H3K27me3 binding by CBX8 while increasing interactions with nucleic acids and participation in variant PRC1. We show that treatment with UNC7040 leads to efficient PRC1 chromatin eviction, loss of silencing and reduced proliferation across different cancer cell lines. Our discovery and characterization of UNC7040 not only revealed the most cellularly potent CBX8-specific chemical probe to date, but also corroborates a mechanism of polycomb regulation by non-histone lysine methylated interaction partners.

## INTRODUCTION

Precise inheritance of distinct gene expression programs during cell division is essential for cellular differentiation and for development of multicellular organisms with stable tissues. Epigenetic mechanisms promote heritable transmission of gene expression and cell identity in the absence of the initial stimulus and without changes in DNA (Moazed, 2011; Reinberg and Vales, 2018). Feedback loops of writing and reading of posttranslational histone modifications contribute to *cis*-acting epigenetic mechanisms. Important regulators of this pathway include Polycomb group (PcG) proteins, which assemble into large multi-subunit complexes that mediate chromatin modifications to enforce transcriptional gene silencing (Aranda et al., 2015; Schuettengruber et al., 2017; Simon and Kingston, 2013). Mutations in PcG protein-encoding genes are frequently associated with malignancies, underscoring the importance of the polycomb pathway in maintenance of cell identity (Chan and Morey, 2019; Laugesen and Helin, 2014). The two major complexes, Polycomb Repressive Complex 1 (PRC1) and PRC2, harbor distinct writing and reading activities of histone modifications. PRC1 complexes are defined by an E3 ligase activity catalyzing monoubiquitination of histone H2A at lysine 119 (H2AK119ub1) (Cao et al., 2005; de Napoles et al., 2004; McGinty et al., 2014; Wang et al., 2004a), while PRC2 complexes generate mono-, di- or tri-methylation of histone H3 at lysine 27 (H3K27me1, 2, 3) (Cao et al., 2005; Czermin et al., 2002; Müller et al., 2002). PRC1 can be further subdivided into canonical and variant complexes (cPRC1 and vPRC1, respectively), based on incorporation of chromobox domain–containing (CBX) proteins. CBX proteins can read H3K27me3 and facilitate PRC1 recruitment to PRC2 target loci (Cao et al., 2002; Min et al., 2003; Wang et al., 2004b). vPRC1 complexes interact with RYBP or YAF2 in place of CBX proteins and localize to chromatin independently of H3K27me3 involving at least in part recognition of unmethylated CpG dinucleotides (Blackledge et al., 2014; Gao et al., 2012; Tavares et al., 2012). The mode of PRC2 recruitment to chromatin remains the subject of intense investigation but some accessory subunits can recognize PRC1-mediated H2AK119ub1 (Cooper et al., 2014; Kalb et al., 2014; Kasinath et al., 2021). Hence, through reciprocal interaction with *cis*-acting histone modifications, cPRC1 and PRC2 engage in a positive feedback mechanism which can sustain heritable gene silencing (Moussa et al., 2019).

Many PcG proteins are represented by multiple paralogs (Hauri et al., 2016; Kloet et al., 2016). Mammalian genomes encode five polycomb-associated CBX proteins (CBX2, 4, 6, 7 and 8) which contribute unique and non-overlapping functions to cPRC1 target selectivity and activity during development and in disease (Chan and Morey, 2019; Laugesen and Helin, 2014). In embryonic and adult stem cells, cPRC1 is mostly associated with CBX7 which promotes self-renewal by silencing differentiation-specific genes including CBX2, 4 and 8. Conversely, CBX2, 4 and 8 are expressed during differentiation and repress genes involved in stem cell maintenance (Creppe et al., 2014; Morey et al., 2013). CBX7 and CBX8 have been linked to cancer through PRC1-dependent transcriptional repression of the well-known tumor-suppressor locus Cdkn2a (also known as Ink4a/Arf) (Dietrich et al., 2012; Gil et al., 2004; Scott et al., 2007; Tan et al., 2011; Yap et al., 2010). In addition, there is evidence in breast cancer and leukemia that CBX8 exerts oncogenic activity through non-canonical functions independent of PRC1 which are poorly defined (Chung et al., 2016; Tan et al., 2011). Our current understanding of how different CBX proteins contribute to PRC1 targeting to chromatin is incomplete. While *in vitro* experiments revealed a range of low specificities and affinities for H3K27me3, *in vivo* studies suggest that chromatin binding is governed by CBX proteins engaging in multiple, distinctive interactions with histone modifications and non-specific binding to DNA/RNA (Connelly et al., 2018; Zhen et al., 2016).

Polycomb’s direct role in transcriptional repression, and misregulation in many disease states (Chan and Morey, 2019; Koppens and van Lohuizen, 2016; Pasini and Di Croce, 2016), has inspired the development of chemical probes to explore polycomb biology, and consequential drug discovery opportunities leading to PRC2-directed inhibitors entering clinical trials (He et al., 2017; Knutson et al., 2014; Knutson et al., 2013; Qi et al., 2017). PRC1 chemical probe development has not yet led to clinical studies, but we and others have worked toward potent ligands for the readers of PRC1, CBX chromodomains (CDs) (Lamb et al., 2019; Milosevich et al., 2016; Ren et al., 2015; Ren et al., 2016; Simhadri et al., 2014; Stuckey et al., 2016a; Stuckey et al., 2016b; Wang et al., 2020). Antagonists of methyl-lysine (Kme) reader domains, such as CBX domains, target the conserved aromatic cage that recognizes methylated lysine via cation-π and hydrophobic interactions (James and Frye, 2016). While to date, all potent ligands for CBX CDs (*Kd* < 1μM) are peptidomimetics, low affinity small molecules have been reported with hints of on-target activity in some assays (Ren et al., 2015; Ren et al., 2016). The most interesting of these, MS351, implicated an allosteric interaction between the aromatic cage of CBX7 and adjacent nucleotide binding residues (Ren et al., 2015). Nucleic acid binding to a broader range of Kme reader domains has been recently reviewed (Weaver et al., 2018), and polycomb CBX CDs have been reported by multiple groups to bind nucleic acids in a non-sequence specific fashion (Connelly et al., 2018; Yap et al., 2010; Zhen et al., 2016).

We have been intrigued by the possibility for allosteric interactions between Kme recognition and nucleotide binding to both regulate endogenous PRC1 activity and influence the cellular activity of CBX by targeted chemical probes. We recently reported a potent cellular positive allosteric modulator (PAM), UNC4976, that binds to CBX7 competitively with H3K27me3 peptides, while increasing affinity for oligonucleotides (Lamb et al., 2019). Both DNA and Kme binding are also critical for chromatin association of CBX8. Yet, how these targeting activities contribute to canonical and non-canonical CBX8 functions in development and disease remains largely unclear. Herein we report the discovery and characterization of UNC7040, a potent, cellularly active PAM specific for CBX8. UNC7040 efficiently impairs cPRC1 targeting and transcriptional repression by disrupting CBX8 interaction with H3K27me3. Notably, the PAM activity of UNC7040 increases CBX8’s affinity for DNA and RNA, redistributes PRC1 to non- H3K27me3 loci, and enhances cellular efficacy to provide a powerful tool to dissect its canonical and non-canonical roles in gene regulation. Together, these results further establish the importance of allostery in CBX cellular pharmacology and activity in disease model systems dependent upon CBX8.

## RESULTS AND DISCUSSION

### Structure-based design of CBX8 selective compounds

During the development of UNC3866, a peptidomimetic cellular probe that selectively targets CBX4/7 CDs (Stuckey et al., 2016a), we observed that the replacement of an alanine methyl group by an ethyl group within UNC3866 modestly decreased the affinity towards CBX7 (∼6-fold), while slightly increasing its affinity toward CBX8 (**Figure 1A,** UNC4939) (Stuckey, 2016; Stuckey et al., 2016a). Moreover, replacement of the alanine residue with an isopropyl group resulted in a compound (UNC4030) that was a 100-fold less potent antagonist of CBX7 and a 2- fold more potent antagonist of CBX8 when compared to UNC3866. Additionally, we recently developed a positive allosteric modulator (PAM) of CBX7, UNC4976, which contains a norbornyl, methyl substitution at the lysine mimetic position instead of the diethyl group present in UNC3866, which binds to both CBX7 and CBX8 to a modestly greater extent *in vitro* and additionally possesses much better cellular efficacy than UNC3866 (**Figure 1A,** UNC4976) (Lamb et al., 2019). Based on these results, we hypothesized that the combination of a more sterically demanding group at the alanine position of UNC3866 with lysine mimetics that could induce positive allostery with nucleotide binding, as seen with UNC4976 (Connelly et al., 2018; Lamb et al., 2019), could result in a potent and selective PAM for CBX8.

**Figure 1.**
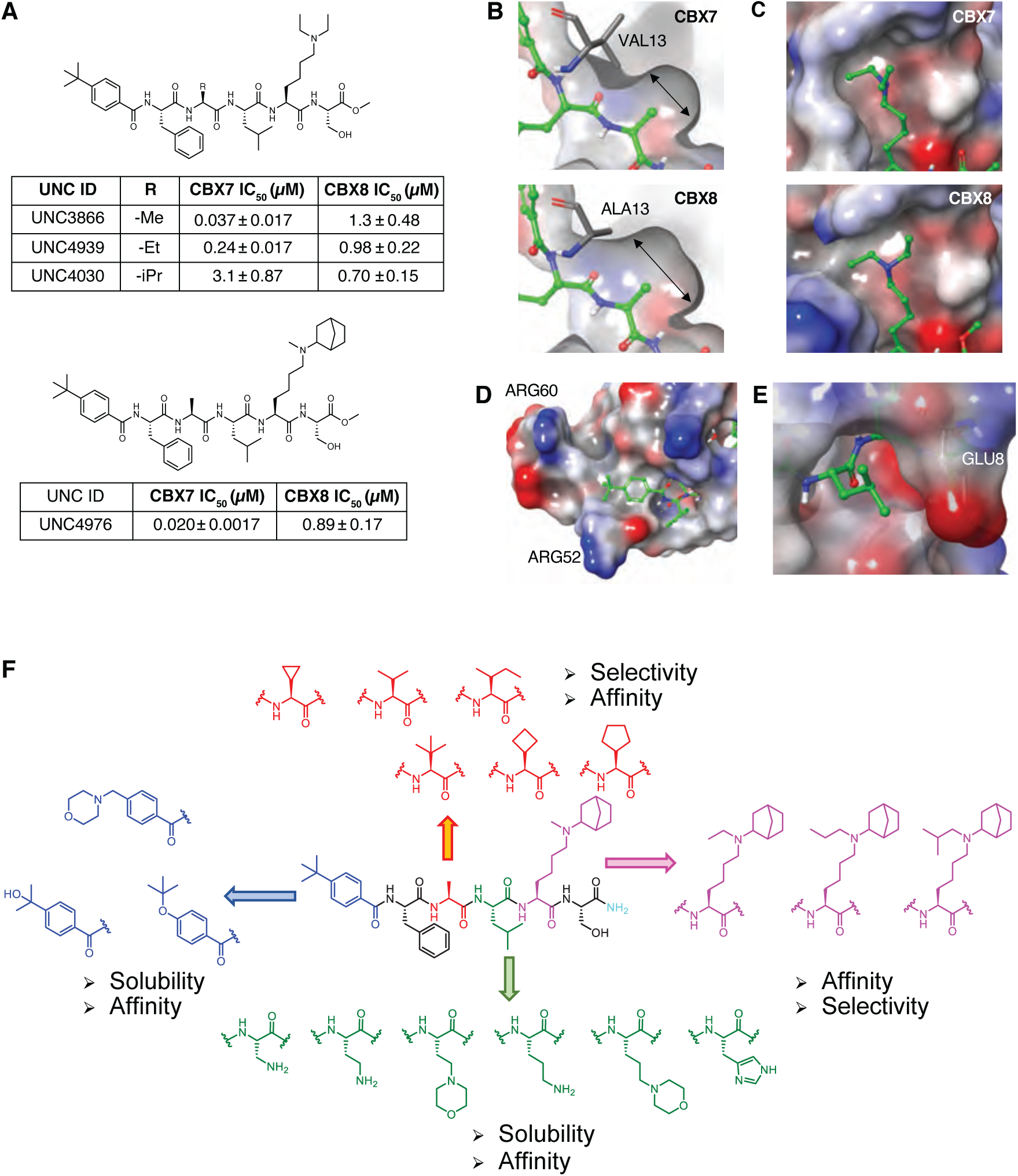
Previously reported CBX7 peptidomimetic antagonists, the structural basis for the design of CBX7 and CBX8 compounds, and SAR strategy using UNC4976 as a template. (A) Previously reported CBX7 compounds and their *in vitro* potency in an AlphaScreen competition assay. (B) Comparison of the alanine binding pocket size of CBX7 (PDB ID: 5EPJ, top) and CBX8 (PDB ID: 5EQ0, bottom). (C) Comparison of the aromatic cage region of CBX7 (top) and CBX8 (bottom). (D) The binding mode of the N-terminal *tert*-butyl benzoyl cap of UNC3866 in CBX8. (E) The binding mode of the UNC3866 leucine side chain in CBX8. UNC3866 is displayed in ball-and-stick with carbons colored green. (F) SAR strategy for new compounds (Red: Ala-position; Blue: N-cap-position; Green: Leu-position; Magenta: Lysine mimetics-position).

In addition to these preliminary structure activity relationship studies, recent efforts to develop CBX7 peptidomimetic antagonists have also provided structural insights into the slight differences between the CBX CDs, CBX2, 4, 7, and 8 (Milosevich et al., 2016; Stuckey et al., 2016a; Stuckey et al., 2016b). Structural analysis of UNC3866 bound to CBX7 (PDB ID: 5EPJ) and CBX8 (PDB ID: 5EQ0) provides a clear rationale for the potency differences between the compounds discussed above (UNC3866, UNC4939, and UNC4030 in **Figure 1A**), wherein the co-crystal structures revealed that CBX7 possesses a much smaller hydrophobic pocket where the side chain of the alanine resides as compared to CBX8 (**Figure 1B**). In the case of CBX7, this small hydrophobic pocket consists of Val13, Val30 and Ile48 whereas the hydrophobic pocket of CBX8 is made up of Ala13, Val30, and Ile48. Consequently, the substitution of Val13 for Ala13 allows CBX8 to accommodate larger functional groups such as an ethyl or isopropyl moiety within this pocket. This difference has also been discussed by Simhadri *et al*. (Simhadri et al., 2014), who observed that the residue at the alanine position is responsible for the selectivity and potency of CBX substrates and peptidomimetic antagonists. Accordingly, we decided to take advantage of this larger binding pocket and other structural differences to aid the design of more potent and selective CBX8 antagonists. Our overall structure activity relationship (SAR) strategy is depicted in **Figure 1F**. Initially, we introduced 6 bulkier and structurally diverse functional groups at the alanine position to investigate the optimum substituent size to occupy this pocket (**Figure 1F**, red color code). In addition to the alanine pocket being a key factor for CBX8 selectivity versus CBX7, the aromatic cage of CBX8 that binds methylated lysine also contains structural differences that could lead to increased selectivity. The UNC3866 co-crystal structure revealed that the aromatic cage of CBX8 is more expansive than the CBX7 aromatic cage (**Figure 1C**). Therefore, introduction of larger functional groups than the methyl group at the lysine residue in UNC4976 could increase the selectivity and potency for CBX8 (**Figure 1F**, magenta color code). Additionally, a limitation of UNC4976 is its diminished water solubility due to the hydrophobic norbornyl group at the lysine mimetic position. Therefore, given that the CBX8 potency and selectivity SAR pointed to even further increases in hydrophobicity, it was apparent that adding a solubilizing group would be necessary to enable biological evaluation while avoiding biologically incompatible solvents, such as >1% DMSO. According to the co-crystal structure, the N-terminal *tert*-butyl benzoyl residue (N-cap) of UNC3866 is solvent accessible and there are several proximal hydrogen-bond donors (HBD) in CBX8 (**Figure 1D**). Therefore, we explored introduction of heteroatoms and protonatable groups at this position, as hydrogen-bond acceptors (HBA) at the N-cap position might be beneficial to potency and solubility (**Figure 1F**, blue color code). Additionally, the side chain of leucine in the UNC3866 co-crystal structure with CBX8 is also solvent accessible, which provides another position to enhance solubility (**Figure 1E**). The co-crystal structure of the H3 peptide (Ren et al., 2015) where this leucine residue is an arginine, also suggests an interaction of this side chain with Glu8 in CBX8. We hypothesized that installation of a HBD at this position could create a favorable interaction with Glu8 to increase the binding affinity and/or solubility of CBX8 ligands (**Figure 1F**, green color code). Because we were interested in tracking SAR trends in multiple assays where trade-offs between in vitro affinity and selectivity, cellular efficacy, and solubility were likely to be required, we decided on an iterative strategy of rounds of synthesis and testing followed by rounds of mixing and matching of optimal substituents, rather than a fully combinatorial approach that would result in synthesis of many mismatched substituents (i.e. combinations of all hydrophobic residues or all polar residues).

### SAR investigation of “the Ala pocket”

Based on our structural analysis described above, we first investigated the effect of replacing the alanine methyl group in UNC4976 with bulkier groups, the key position to achieve CBX8 selectivity. We synthesized peptidomimetic compounds using conventional solid-phase peptide synthesis with Rink-Amide Resin and tested their affinity to CBX7/8-CD using a time-resolved fluorescence energy transfer (TR-FRET) assay (Rectenwald et al., 2019). As expected, CBX8 could accommodate larger substituents at this position relative to CBX7 (**Table S1**). Installation of an isopropyl group (**2**) increased selectivity for CBX8 200-fold compared to a methyl group (**1**). When the alanine was replaced with an isoleucine (**3**), the compound lost all potency at CBX7 (> 30 μM, **Table S1**), which increased the CBX8 selectivity ∼54-fold, but reduced overall CBX8 affinity compared to the isopropyl-containing compound (**2**). Both CDs were able to tolerate the cyclopropyl group to generate potent CBX7 and CBX8 antagonists albeit with less than 2-fold selectivity (**4**). The use of a cyclobutyl group (**5**) or cyclopentyl group (**6**) retained CBX8 affinity effectively while reducing CBX7 affinity more than 6-fold compared to the isopropyl moiety (**2**). The addition of a *tert*-butyl group (**7**) was unfavorable for binding at both CBX8 and CBX7, indicating that this side chain size was too large for either pocket. Compound **2**, which has valine at the alanine position, was a potent antagonist for CBX8 with 21-fold selectivity over CBX7, providing an ideal scaffold to explore SAR further at the N-cap position.

### SAR investigation of “the N-cap”

Next, we focused on the effects of N-cap modifications on solubility and CBX8 binding affinity. We modified **2** to contain an isopropanol (**8**), morpholino (**9**), and *tert*-butoxy group (**10**) and found that, interestingly, the introduction of just one heteroatom at the N-cap position results in full water solubility at 10 mM, which is not achievable with UNC4976. Overall, there were small differences in CBX8 potency between these 4 variations (**Table S1**). The isopropanol group (**8**) did not have improved selectivity against CBX7 compared to **2**. Slight differences between the morpholino group (**9**) or *tert*-butoxy group (**10**) decreased the affinity to CBX7 approximately 2- fold, which modestly increased their CBX8 selectivity. However, we observed cleavage of the *tert*- butoxy group to the hydroxyl group during HPLC purification (0.1% TFA) and synthesizing **8**-**10** required two more synthetic steps compared to the synthesis of compound **2** making the route less attractive. Hence, we decided to retain a *tert*-butyl substitution at this position and rely on introduction of a solubilizing group at the leucine position of UNC4976, which is an alternative solvent accessible site.

### SAR investigation of “the Leu position”

Once again, we explored the leucine position starting with compound **2** as our scaffold, based on CBX8 affinity and selectivity. Analogues containing hydrogen-bond donors with differing linkers from the peptide backbone at the leucine position were synthesized (**Table S1**). Similar to the N-cap modifications, adding one or more heteroatoms at this position was highly advantageous for the compound’s water solubility. Introduction of an amino group with a distance of two-carbon atoms from the backbone carbon (**11)** slightly decreased potency for both CBX8 and CBX7, whereas compound **13** which has an amino group with a distance of three-carbon atoms showed modestly increased potency for CBX8 and selectivity over CBX7 (**Table S1**). Morpholino substitution with either a distance of two or three-carbon atoms as in compound **12** and **14**, respectively, slightly increased the potency toward CBX8 while decreasing potency toward CBX7, overall increasing CBX8 selectivity to 24 to 58-fold. We also prepared compound **15** containing a histidine side chain, which we expected to be able to form a hydrogen bond with the side chain of Glu8 in CBX8 (**Figure 1E**). However, histidine substitution caused complete loss of CBX7 potency as well as dramatically reduced CBX8 potency. Taken together, these data indicate that introducing protonatable amines at the leucine position can improve compound solubility and CBX8 selectivity. Compound **14** containing a morpholino group with a three-carbon atom linker showed the best improvement in CBX8 potency and selectivity among all these variations, and we selected this substituent to move forward with additional SAR investigations. The pKa of morpholine is also in a more favorable range for cell penetration as compared to the more basic primary amines of compounds **11** and **13**.

### SAR investigation of “Lys mimetics”

Finally, we evaluated the effect of incorporating a group bulkier than the methyl in UNC4976 at the lysine mimetic position. We maintained the norbornyl substitution from UNC4976, as this group had been shown to convey PAM activity in the interaction of oligonucleotides with CBX7, resulting in greatly enhanced cellular efficacy (Lamb et al., 2019). To investigate modifications at this position, we combined the optimal substituents defined above: an isopropyl (**2**) or cyclobutyl (**5**) group at the alanine position and the 3-carbon atom spaced morpholino group at the leucine position (**Table S1**).

Accordingly, we designed and synthesized 8 compounds which contained either ethyl, propyl, or isobutyl substitution on the norbornyl-lysine mimetic (**Table S1**). Ethyl or isobutyl substitution in compounds **16** and **20** and compound **18** and **22**, respectively, increased potency for CBX8 approximately 2-fold, but also increased potency for CBX7, resulting in similar selectivity. Installation of the propyl group (**17** and **21**) showed negligible effect on both CBX8 and CBX7 potency. Overall, incorporation of a larger alkyl group did not result in any significant enhancement in CBX8 *in vitro* potency and selectivity compared to the compounds containing a methyl group (**14** and **19**). However, as discussed below, variation at this position had a dramatic effect on cellular activity.

### Cellular screen of new antagonists of CBX8

Previous studies describing PAMs of CBX7 revealed that UNC4976 (**Figure 1A**) demonstrates 14-fold enhanced cellular potency compared to UNC3866 (**Figure 1A** top compound), despite almost identical *in vitro* thermodynamic and kinetic affinity profiles, as well as cell permeability (Lamb et al., 2019; Moussa et al., 2019; Stuckey et al., 2016a; Stuckey et al., 2016b). Based on this finding, we evaluated our compounds in a CBX8-specific cellular reporter assay to determine if cellular potency trends differ substantially from *in vitro* results. To evaluate cellular efficacy in the context of native chromatin, we engineered a mouse embryonic stem cell (mESC) line that expresses a CBX8 protein fused to the Tet repressor domain (TetR). This assay is similar to the recently developed Green Fluorescent Protein (GFP) reporter CBX7 cellular assay (Moussa et al., 2019), however this mESC line contains a landing site with a cassette that harbors an artificial DNA binding array composed of 12xZFHD1, 4xGAL4 UAS, and 7xTetO sites upstream of a PGK promoter controlling the expression of Puromycin resistance and a GFP gene (**Figure 2A**). The TetR-CBX8 fusion enables recruitment and assembly of a functional canonical PRC1 (cPRC1) complex at a TetR-DNA binding site upstream of an active GFP reporter gene (**Figure S1A, B)**. After this nucleation event, additional PRC1 and PRC2 complexes are recruited to sites along the gene body through interactions of the CBX8 CD with trimethylated histone H3, lysine 27 (H3K27me3), which results in formation of a repressive polycomb chromatin domain similar to endogenous targets and transcriptional silencing of the GFP gene as measured by a decrease in GFP signal using flow cytometry (**Figure 2B, Figure S1B, C**). Notably, GFP reporter gene repression is reversable upon Doxycycline (DOX)-dependent release of the TetR fusion protein (**Figure S1D**). Thus, while ectopic targeting of CBX8 can initiate formation of a repressive polycomb chromatin domain in mESCs, in contrast to CBX7 it fails to promote propagation of silencing in the absence of the initial stimulus (Moussa et al., 2019). Having established this reversible CBX8 reporter, we reasoned that addition of CBX8 antagonists would inhibit the binding of endogenous PRC1 mediated by CBX8, prevent assembly of additional cPRC1 complexes along the gene body, and de-repress the GFP reporter gene.

**Figure 2.**
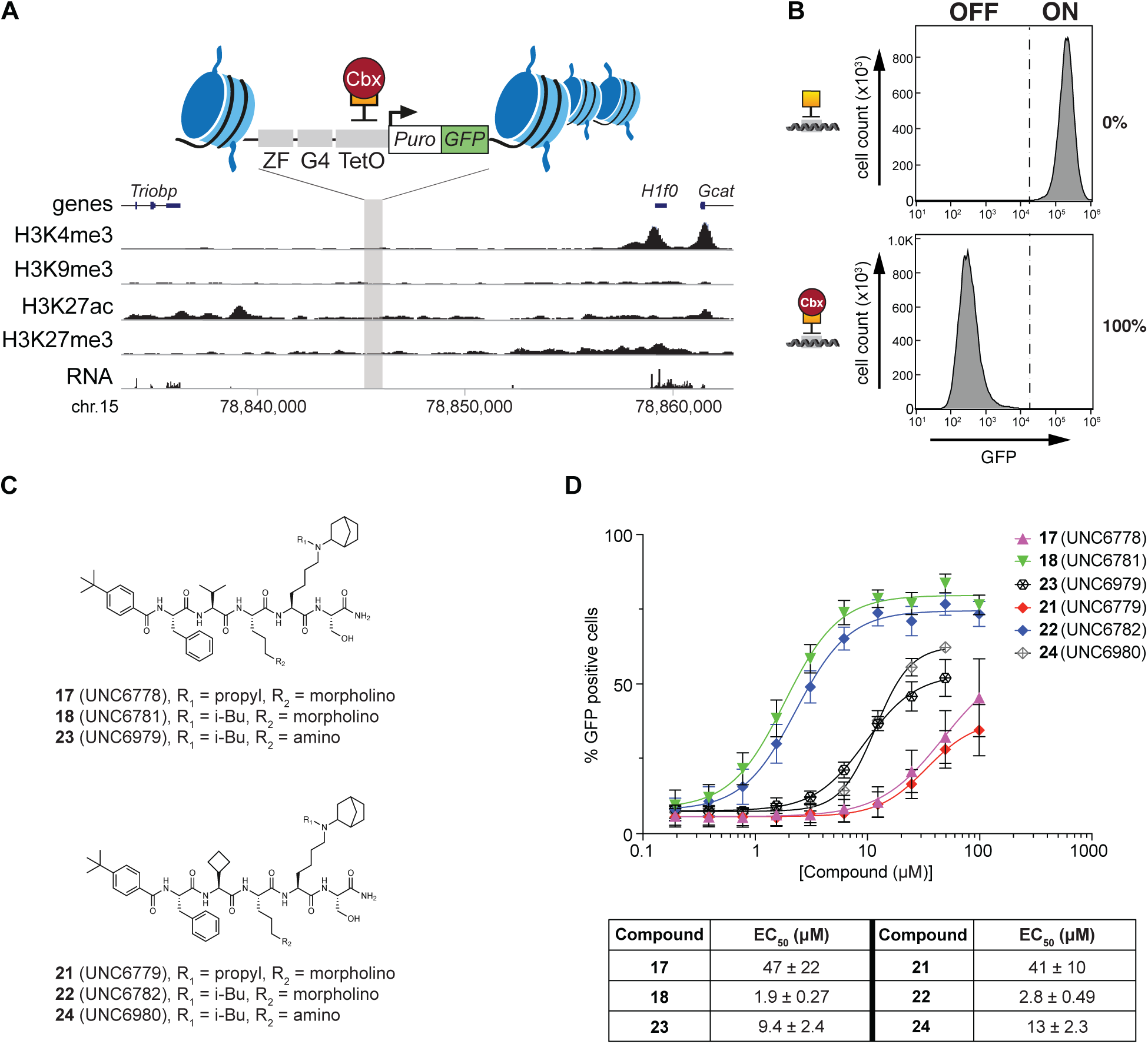
Design and readout of the CBX8 GFP reporter assay and optimized compounds from SAR studies and their cellular efficacy. (A) Schematic representation of the CBX8- specific GFP reporter gene construct containing an array of DNA binding sites (12XZFHD1, 4XGAL4 UAS, 7XTetO) located upstream of a constitutive PGK promoter driving expression of a Puromycin-IRES-GFP reporter gene. TetO DNA binding sites facilitate recruitment of TetR fused to human CBX8. (B) Flow cytometry histogram shows % repression of GFP expression in response to recruitment of TetR alone (top) or TetR-CBX8 (bottom) in the absence of CBX8 antagonists. (C) Compounds with the best potency profile from the SAR studies. (D) The CBX8 reporter assay results of the 6 optimized compounds. Data shown are mean ± SD, n = 9.

Although our SAR studies led to some highly water soluble CBX8 antagonists, we performed a DMSO tolerance test using a Celltiter-Glo cell viability assay in the mESC line utilized in our CBX8 GFP reporter assay so that less soluble analogues could also be assessed at non-toxic DMSO concentrations. We found that treatment with up to 1.25 % DMSO was tolerated in our mESC cell line with no cellular toxicity effects observed (**Figure S1E**). Accordingly, we assessed the cellular efficacy of our CBX8 antagonists discussed above with final assay concentrations of 1% DMSO. After 48-hours exposure of test compounds in the CBX8 reporter mESC line, we determined the change of GFP levels compared to 1% DMSO control by flow cytometry. Results organized as discussed above for our *in vitro* SAR are depicted in **Figure S2A- D**.

Consistent with CBX8 potency trends in our TR-FRET results, compounds with alanine position substituents isopropyl (**2**) or cyclopropyl (**4**) showed the largest reactivation of GFP signals among the 7 compounds, followed by cyclobutyl (**5**) and cyclopentyl (**6**) groups, while the methyl (**1**), isobutyl (**3**), and *tert*-butyl (**7**) all demonstrated minimal effect on GFP reactivation (**Figure S2A**). Overall, none of the modifications made at the alanine position fully reactivated the GFP signal. Next, we examined the N-cap modifications. Although there were no significant *in vitro* potency differences between these four N-cap variations (compounds **2**, **8**, **9**, and **10**), cellular assay data showed that the *tert*-butyl moiety from the parent compounds, UNC3866 and UNC4976, was the most potent modification (**Figure S2B**). For the Leu position modifications, ornithine substitution (**13**) at this position showed the highest GFP reactivation compared to morpholino (**12** and **14**) or leucine (**2**) containing compounds at the highest tested concentration (99 μM) (**Figure S2C**). Previous studies with cell penetrating peptides (CPPs) reported that some CPPs, which usually contain a positively charged portion in the molecule, may be able to reach a high enough local concentration on the cell membrane to form peptide-rich domains that can transiently disrupt the membrane or promote peptide translocation into the cell (Duchardt et al., 2007; Herce et al., 2009; Tunnemann et al., 2006). Because the positive charge of the ornithine substitution is the most solvent exposed among the five tested compounds, this transient plasma membrane permeabilization theory may potentially explain why this particular molecule showed the best activation but only at the highest concentration of compound tested. Lastly, lysine mimetic modification with a bulkier group than the methyl present in UNC4976 at this position had a dramatic effect on cellular efficacy in contrast to the minimal efficacy observed with modifications at the other amino acid sites (**Figure S2D**). Surprisingly, although none of these lysine mimetic variations caused a significant boost in CBX8 *in vitro* potency (**Table S1**), cellular efficacy was dramatically changed. In general, longer chained and bulkier groups at the lysine mimetic position displayed increased cellular efficacy. The compounds with isobutyl groups, **18** and **22**, demonstrated the best GFP activation with almost fully reactivated GFP levels in contrast to the other compounds in the series, which did not reactivate GFP by greater than 50%. This striking result showed that increased steric bulk at this position that did not affect *in vitro* potency influenced cellular efficacy to a large extent.

Since the ornithine substitution (**13, Figure S2C**) at the leucine position and the isobutyl group at the lysine mimetic position (**18, Figure S2D**) showed the greatest cellular efficacy, we incorporated these modifications into one molecule and synthesized two additional compounds that mix the best substituents at each position (**Figure 2C**, compound **23** and **24**). The four best compounds from the previous SAR study and these additional compounds **23** and **24** were then tested in the CBX8 cellular assay to compare the effect of combined optimal modifications. While *in vitro* results (**Table S1**, compound **18** vs **22**) demonstrated a more than 3-fold *in vitro* potency difference between isopropyl and cyclobutyl groups at the alanine position, the cellular potency difference was less than 2-fold (**Figure 2D**, **17** vs **21**, **18** vs **22**, and **23** vs **24**). This data shows that the isobutyl group at the lysine mimetic position is more influential for cellular potency than the isopropyl or cyclobutyl side chain of the alanine position modification. Although we expected **23** and **24** to be more efficacious antagonists in the cellular assay than **18** and **22** because the ornithine-containing compound (**13**) displayed the strongest GFP activation from the SAR study, the morpholino containing compounds demonstrated 5-fold more potent EC_50_’s (EC_50_ of **23** = 9.4 ± 2.4 µM and EC_50_ of **24** = 13 ± 2.3 µM versus EC_50_ of **18** = 1.9 ± 0.27 µM and EC_50_ of **22** = 2.8 ± 0.5 µM). Interestingly, isobutyl containing compounds, **18** and **22,** displayed 20–30-fold enhanced cellular potency compared to the propyl containing compounds, **17** and **21**. We hypothesized that this could be due to enhanced allosteric influence of the isobutyl compounds on CBX8 nucleic acid binding, similar to the allostery we observed with UNC4976 versus UNC3866 with CBX7 (Lamb et al., 2019). An allosteric mechanism is plausible based on a recent NMR study of CBX8, which revealed that trimethylated H3K27 binds to one face of CBX8 while DNA binds to the other face with the aromatic cage where our lysine mimics bind sandwiched in between (Connelly et al., 2018). However, before seeking other explanations for the dramatic difference in cell potency mediated by addition of a single methyl group to convert the propyl substituent in **21** to an isobutyl group in **22**, we examined the cell permeability of our compounds, as this could also be a significant variable influencing cell potency.

### Influence of modifications on cellular permeability

An important factor to consider with these compounds is how cell permeability changes with each modification as this can influence cellular efficacy. To address this question, we utilized the chloroalkane penetration assay (CAPA) (Peraro et al., 2018; Peraro et al., 2017) to quantitatively measure cell permeability. CAPA is especially useful in rank ordering compounds of relatively low permeability that cannot be rank ordered by traditional permeability assays, such as Caco-2 or PAMPA, because of inadequate dynamic range at the lower limits of detection (Foley et al., 2020). In order to systematically compare the effect of each functional group modification on cell permeability, we synthesized 9 compounds containing a chloroalkane tag as depicted in **Table S2**.

Assessment of CT-modified compounds by CAPA revealed that incorporation of a cyclobutyl group at the alanine position had little effect on the permeability of the compound compared to the parent compound with almost equipotent CP50 values (**Table S2**, **25** vs **26** and **Figure S2E**). For the N-cap position the *tert*-butyl was the most permeable, which correlates with the potency trend observed in the CBX8 reporter assay (**Table S2**, **26** and **28** and **Figure S2F**). However, based on their CP50’s, the greatly diminished cellular activity for the compounds containing a hydroxy (**27**) or morpholino (**28**) functional group at the N-cap position are not fully attributable to permeability (or *in vitro* affinity (**Table S1**)) and there could be other factors beyond cell permeability that influence their cellular activity. At the leucine position, the leucine functional group increased permeability relative to compounds containing a positively charged side chain (**Table S2**, **26** vs **29-30** and **Figure S2G**). Interestingly, ornithine-containing compound (**30**) demonstrates a very steep CP50 curve above a certain concentration (between 11 and 33 μM), which is similar to our data the CBX8 reporter assay (**Figure S2C**). This is consistent with the aforementioned transient plasma membrane permeabilization (Duchardt et al., 2007; Herce et al., 2009; Tunnemann et al., 2006). Critically, variations at the lysine mimetic position demonstrated no differences in cell permeability (**Table S2**, **30-33** and **Figure S2H**). Therefore, although permeability data explains the increased cell efficacy observed between compounds with leucine position modifications, neither cell permeability nor *in vitro* affinity can explain the significant difference in cellular efficacy of compounds **21** and **22** that differ only at the lysine mimetic position.

### Comparison of 21 (UNC6779) and 22 (UNC6782) by electrophoretic mobility shift assays

The ability for the isobutyl lysine functionalized compounds to more effectively disrupt PRC1 activity in cells as compared to propyl substitution, and the similar permeability and *in vitro* profile of these compounds, encouraged us to investigate the underlying mechanism for this difference by testing the impact of compound **21** and **22** on DNA’s ability to bind to CBX8. While interaction with RNA may also be relevant for CBX CDs (Lamb et al., 2019), DNA has been validated as a contributor to the binding of CBX8 to nucleosomal DNA by NMR and biophysical studies, and its relative stability is advantageous experimentally (Connelly et al., 2018). We first utilized electrophoretic mobility shift assays (EMSAs) to examine whether **21** and **22** stabilize or destabilize the interaction between CBX8 and fluorescently tagged double stranded DNA (FAM-dsDNA). CBX8 CD titration with a constant FAM-dsDNA concentration of 500 nM indicated that at concentrations above 31.3 µM protein we could observe formation of a CBX8-DNA binary complex (**Figure S3A**). A recent study with UNC4976 suggested that the compound’s enhanced cellular efficacy was mainly derived from the ability to stabilize nucleic acid engagement with CBX7 in a ternary complex with UNC4976 acting as a PAM of nucleotide binding (Lamb et al., 2019). We hypothesized that if this mechanism was occurring with CBX8, we would be able to detect ternary complex formation with a protein concentration below that required for ds- DNA/CBX8 association (**Figure S3A)**. Hence, we carried out a compound titration experiment with 15 µM protein concentration. Interestingly, and also consistent with the previous results with UNC4976, treatment with compounds stabilized the CBX8-DNA interaction as observed by the appearance of a fluorescent band **(Figure S3B**, **left**). Increasing concentrations of **22** generated stronger fluorescent bands than similar concentrations of **21** suggesting that the presence of an isobutyl group in the Kme mimetic stabilized the CBX8-DNA complex to a greater extent than a propyl group. The Coomassie stain also suggested enhanced CBX8-DNA binding based on the diminished amount of DNA-free protein with increasing concentrations of **22** (**Figure S3B**, **right**). Taken together, our EMSA data reinforces the hypothesis that **22** possesses an improved ability to stabilize the binding of CBX8 and DNA relative to **21**, but other techniques were clearly required due to the qualitative nature of this assay. Unfortunately, we were not able to utilize fluorescence polarization (FP) to quantitatively examine ternary complex formation (ligand:CBX8:ds-DNA), in contrast to our results with CBX7 (Lamb et al., 2019). Therefore, we approached ternary complex characterization with NMR and molecular dynamics.

### NMR studies of CBX8 with 21 (UNC6779), 22 (UNC6782) and DNA

In order to provide structural insights into the interaction between our ligands, CBX8 and DNA we utilized nuclear magnetic resonance (NMR) spectroscopy. ^1^H-^15^N-HSQC titrations were performed by collecting spectra of the ^15^N-CBX8-CD in the apo state and upon addition of **21** or**22**. Addition of increasing concentrations of **21** and **22** led to significant chemical shift perturbations (CSPs) in the CBX8-CD spectrum consistent with an interaction between the inhibitors and CBX8-CD (**Figure S4A**). Additionally, several residues disappeared upon inhibitor binding which is consistent with these regions undergoing a conformational exchange on an intermediate time scale. Interestingly, several minor peaks appear only in the **22** titration, which are not seen in the **21** titration, indicating that the CD can adopt a minor population structural state when bound to **22** but not **21**. As a distinct set of peaks is seen, this indicates that the minor population is either a fully stable separate population or that the major and minor population are in slow exchange between each other on the NMR time-scale.

To further investigate the structural basis for the interaction of **21** and **22** with CBX8-CD, a normalized change in chemical shift from the apo to inhibitor bound states was calculated using a set of previously published assignments for the CBX8-CD (Connelly et al., 2018). Importantly, this initial analysis is only for the major state population of the **22**-bound CBX8-CD. Residues with significant CSPs upon addition of **21** and **22** map largely to the aromatic cage region and the N-terminal portion of the αA-helix, while residues that broaden beyond detection are in the N-terminal portion of the β1 strand that is known to undergo a conformational change upon binding H3K27me3 (**Figure S4B, C**). Notably, the CSPs are in good agreement with previous data for the addition of the canonical H3K27me3 ligand (**Figure S4B, C**). The residues with CSPs (14/14) and residues that broaden beyond detection (4/5) in the titrations of **21** and **22** titrations are almost identical, indicating the major population CD binds to the two inhibitors largely in a similar manner. To assess whether the CBX8-CD can interact with **21** or **22** and DNA simultaneously, ^1^H-^15^N-HSQC spectra were collected on the ^15^N-CBX8-CD in the apo state, inhibitor bound state and inhibitor plus DNA bound states. Addition of an 11bp DNA with the CBX8-CD pre-bound to either **21** or **22** led to significant CSPs in the CBX8-CD spectrum indicating binding (**Figure 3A**). Comparison of the CBX8-CD spectrum bound to DNA only, inhibitor only, and inhibitor + DNA reveal the bound states are unique from each other consistent with the formation of a ternary complex with addition of inhibitor + DNA (Weaver et al., 2018). Residues with significant CSPs upon addition of the 11bp DNA to the CD pre-bound to either **21** and **22** (major population) map largely to the C-terminal portion of the β1 strand and αA helix consistent with the previously determined 11bp DNA binding site identified in the absence of inhibitor (**Figure 3B, C**). Comparison of the residues with significant CSPs upon addition of DNA reveal 11 residues that are identical between **21** and **22** binding, which indicates a relatively similar binding mode. However, a small subset of CSPs upon addition of DNA to the inhibitor bound CD were different, these included S36 and Y39 for **21**-bound and K33, E43, L49, A51 and A56 for **22**-bound suggesting there are subtle differences in the structural mechanism of DNA binding for each ligand. Notably, for the **22**-bound CBX8-CD CSPs were also seen in the subset of peaks corresponding to the minor population, indicating that it also associates with DNA. Though the structure of this minor population is currently unknown, it is clearly active with respect to DNA binding.

**Figure 3.**
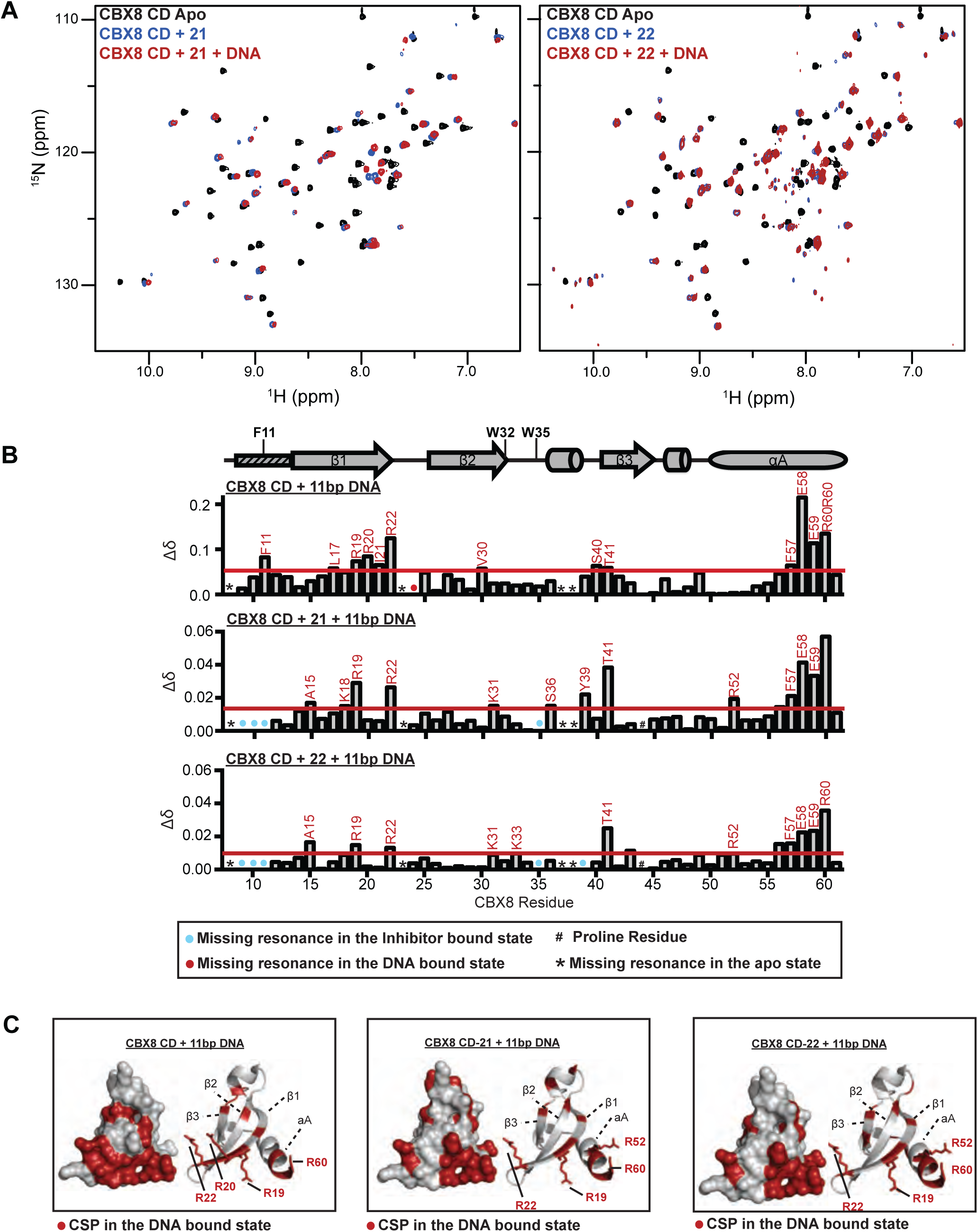
CBX8-CD residues with CSPs upon binding to 11bp DNA only and 11bp DNA with 21 or 22. (A) Full ^1^H-^15^N-HSQC overlays for ^15^N-CD upon addition of compound 21 (left) or 22 (right) in the absence and the presence of 11bp DNA. (B) Normalized CSP (Δδ) between 11bp DNA-bound (top), 11bp DNA and 21-bound (middle), and 11bp DNA and 22-bound (bottom) spectra are plotted against CBX8 residue number. (C) Residues with significant CSPs upon addition of 11bp DNA only, 11bp DNA with compound 21, and 11bp DNA with compound 22 plotted onto a cartoon and surface representation of the CD and colored red.

Taken together, NMR data indicate that both **21** and **22** bind to the CBX8-CD and form a DNA-ternary complex. For the majority of resonances, the chemical shifts upon association with **21** or **22** alone lie along a near linear trajectory towards the tertiary complex chemical shifts, supporting that these compounds stabilize a DNA-binding competent conformation of the CBX8- CD. However, by analyzing the major state bound populations of **21** and **22**, we could not identify significant differences in the inhibitor binding mode. The presence of a minor state bound population of the CBX8-CD with **22** could be the key to the differences we detected for these two inhibitors in a cellular context and we decided to explore this further using molecular dynamics.

### Molecular Dynamics of CBX8 with 21 (UNC6779) and 22 (UNC6782)

In general, an allosteric modulator of a protein acts by altering the host protein’s conformational ensemble which in turn may alter its binding affinity to a third molecule at a remote binding site (a positive value of *α* in the Stockton/Ehlert allosteric binding model) (Boehr et al., 2009). We therefore applied molecular dynamics (MD) simulations to explore how the ligands (**21** and **22**) affect the conformational ensembles of CBX8 and analyzed the respective MD trajectories to infer a structural mechanism by which **22** may allosterically enhance the affinity of CBX8 for DNA/RNA. In this study, a total of ∼50 microseconds (μs) of MD simulations on systems including the CBX8 CD in complex with respectively **21** and **22**, as well as the CBX8 CD alone were performed. Structural snapshots, one per 40 picoseconds (ps), were extracted from the MD trajectories, aligned and subjected to a cluster analysis. The analysis was performed in such a way that each cluster contained closely related protein folds, within ∼1 Å of root mean square distance (RMSD). Hence, a centroid of each cluster approximates a distinct conformation within the protein’s conformational ensemble. Of particular interest were clusters that predominantly consisted of the snapshots featuring either **21** or **22**. Indeed, such clusters can be associated with ligand-induced conformations of the ensemble. As hypothesized, we observed that both **21** and **22**, each in its unique way, alter the conformational ensemble of the CBX8 CD. Of the total of 395 conformations identified in all three simulated systems, 94 and 40 were induced by **21** or **22**, respectively (**Figure 4A** and **B**), which suggests a significantly higher conformational mobility of the protein-bound **21**. These ligand-induced conformations were observed during 24% and 31% of time, respectively, for **21**- and **22**-bound CBX8. The existence of such ligand-induced conformations supports the idea that the enhanced binding of the **22**-CBX8 complex to DNA/RNA might be due to the compound’s ability to induce “DNA/RNA-friendly” CD conformations. Some of these ligand-specific conformations could represent the "minor state" inferred from the NMR data for **22**. Moreover, the clustering data suggest a higher average stability of the conformations induced by **22** (40 conformations account for 31% of the simulation time, compared to 94 conformations / 24% time for **21**). This observed stability is consistent with the NMR spectra showing a slower exchange between the major and minor populations for **22**. A 31% share of such conformations in the ensemble of the **22**-bound CBX8 seems sufficient to make a measurable difference in binding. Moreover, it should be kept in mind that this share may dynamically change in the presence of DNA/RNA since the latter would capture and preserve these DNA/RNA-friendly CBX8 conformations as soon as they are spontaneously produced, shifting the overall conformational equilibrium in favor of these species. While we expected ligand-induced conformations to show more focused changes around the aromatic cage, as the two compounds only differ in the methyl-lysine mimetic that is expected to bind in this region (Stuckey et al., 2016a), compound-induced conformations instead reflected broader, CD-wide shifts. This suggests that although the propyl (**21**) to isobutyl (**22**) change is modest in the context of the entire peptidomimetic scaffold, **22** has the ability to drastically alter the conformation of the CBX8 CD even outside the aromatic cage and thereby allosterically enhance DNA/RNA binding.

**Figure 4.**
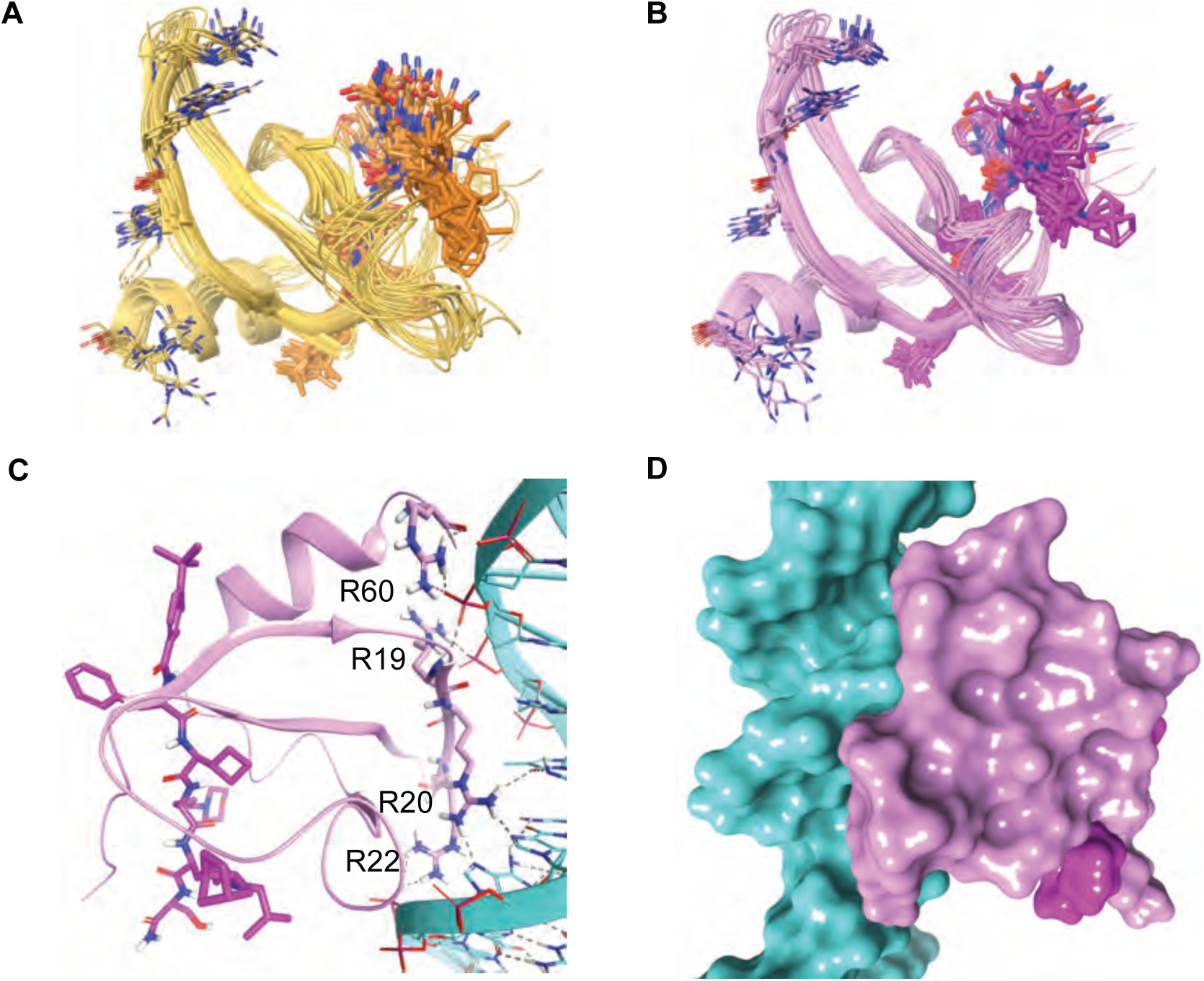
CBX8-Molecular Dynamics (MD) (A) MD simulation snapshots sampled from conformational ensembles of CBX8 in complex with **21** (yellow cartoon / orange sticks) and **22** (pink cartoon / magenta sticks). (B) and (C) An example of a top ranked docking pose showing a large contact surface area between the CBX8-**22** complex (pink and magenta surfaces) and DNA double helix (cyan surface) (B) with the K18-G24 b-strand binding deep into the major groove (C). Only **22**-bound CBX8 (pink cartoon / magenta sticks) has shown binding modes with all four essential DNA-binding residues (R19, R20, R22, and R60) simultaneously interacting with DNA (cyan cartoon/sticks) (C).

We then investigated a possible structural mode of the ligand-induced interaction of CBX8 CD with the DNA double helix. To this end, three sets of 10 MD snapshots each were selected at random from structural clusters predominantly containing either ligand-bound (**21** or **22**) or ligand-free CBX8 CD. All 30 structures were then submitted to automated protein-DNA docking simulations by the High Ambiguity Driven protein-protein DOCKing (HADDOCK) algorithm (van Zundert et al., 2016). The resulting HADDOCK scores were in the range between -70 and -110 kcal/mol that is typical of a small-size protein. Remarkably, HADDOCK scores are consistent with the experimental data, *i.e.*, cluster-weighted averages for CBX8:**22** and CBX7:**21** are, respectively, -81 ± 7 kcal/mol and -73 ± 5 kcal/mol. The top ranked docking poses show a large contact surface area between the protein and DNA (**Figure 4C** and **D**) with the K18-G24 b-strand binding deep into the major groove. Both ligand-bound CDs share significant similarities in the way they bind to DNA. In particular, the protein-DNA interaction implicates the residues R19, R20, R22, and R60 which have been previously identified as important for the interaction with DNA (**Figure 4C**) (Connelly et al., 2018). However, **21**- and **22**-bound CBX8 displayed significant local differences in engaging the DNA double helix. In particular, only **22**-bound protein obtains binding modes with all four essential DNA-binding residues (R19, R20, R22, and R60) simultaneously interacting with DNA (**Figure 4B** and **C**). Moreover, the latter binding modes were adopted by the most populated ligand-induced conformation clusters wherein the cluster-weighted averages of the number of essential residues simultaneously bound to DNA were 2.4 ± 0.3 and 3.1 ± 0.3 for respectively **21**- and **22**-bound CBX8 CDs. Overall, the combination of MD clustering and HADDOCK docking data suggests that modification of the Kme mimetic significantly affects the dynamics of both the K18-G24 loop and the aromatic cage, which in turn are implicated in CBX8:DNA binding, thus providing a clear structural rationale for the positive allosteric effect of **22**.

### Final cellular probe optimization, negative control design and characterization

Considering CBX8 affinity, selectivity against CBX7, cellular efficacy, and cell permeability, we selected compound **22** as the basis for further studies. Although they have similar profiles, we selected the cyclobutyl functional group (**22**) over the isobutyl (**21**) at the alanine position, because unnatural amino acids incorporated into peptidomimetics are known to be beneficial for metabolic stability (Blaskovich, 2016; Qvit et al., 2017). While for synthetic ease, our SAR studies and some mechanistic studies were carried out with C-terminal amides, C-terminal methyl esters are known to show higher cellular efficacy due to improved permeability (Lamb et al., 2019; Stuckey et al., 2016a). Consequently, compound **34**, which has a *tert*-butyl benzoyl at the N-cap position, cyclobutyl glycine at the alanine position, morpholino ornithine at the leucine position, norbornyl, isobutyl lysine at the lysine mimetic position, and a methyl ester at the C-terminus, was synthesized as our final CBX8 cellular probe (UNC7040, **Figure 5A**, top). UNC7040 was profiled in the CBX8 reporter assay and the cell titer glow viability assay, which resulted in an EC_50_ of 0.84 ± 0.11 µM (n = 9) and no cytotoxicity up to 100 µM, respectively (**Figure 5B, C**). The cellular potency enhancement of UNC7040 versus **22** (EC_50_ 2.8 ± 0.5 µM) was as expected for replacing a primary amide with a methyl ester (Lamb et al., 2019).

**Figure 5.**
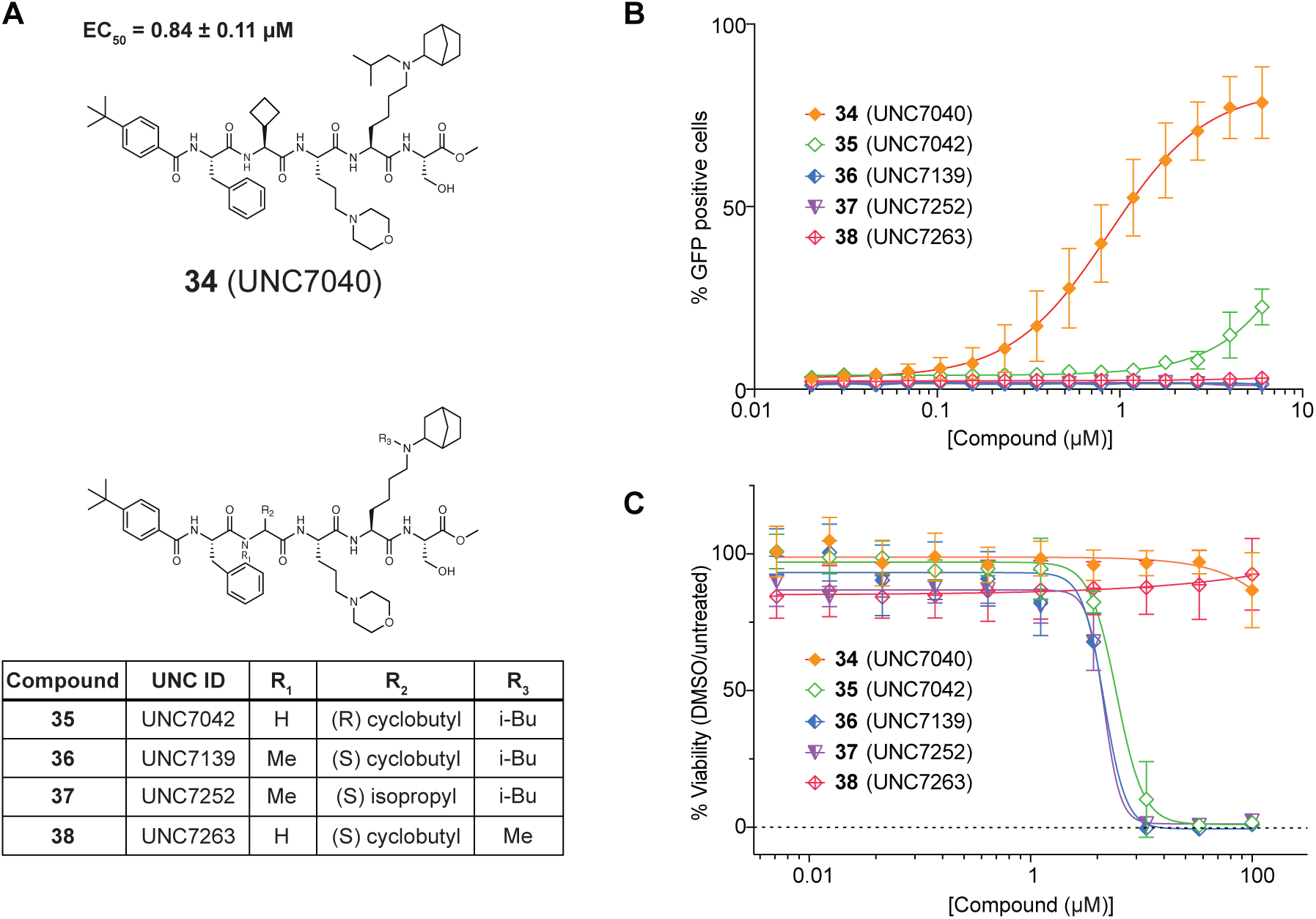
Structures of final CBX8 cellular probe and negative controls and their CBX8 reporter assay and corresponding cell toxicity results. (A) structures of the final probe and the negative controls. (B) CBX8 GFP reporter assay results. (C) Compound cell viability results. n = 9.

We then set out to develop an appropriate negative control compound for UNC7040. The availability of a negative control is important in order to enable cellular and *in vivo* studies to be carried out with a matched pair of compounds that are as similar as possible in their structure, physical properties, and off-target profiles, while differing greatly in their on-target activity. Our search for a negative control for UNC7040 illustrates some of the challenges in creation of the ideal negative control that are perhaps not widely appreciated. Based on prior SAR, that showed that epimers at the leucine position did not bind to CBX CDs (Stuckey et al., 2019), we substituted L-cyclobutyl glycine for D-cyclobutyl glycine resulting in compound **35** (**Figure 5A**, bottom). As expected, compound **35** displayed a negligible effect on de-repression of GFP in the CBX8 reporter assay (**Figure 5B**). However, unexpectedly, this compound possessed much higher cellular toxicity (DC_50_ of 6.0 ± 1.4 µM) than UNC7040 (**Figure 5C**). A negative control compound with cell toxicity not present in the positive control is not a useful comparator, therefore, we explored alternatives. An analogue of the active compound UNC7040 was synthesized with a methylated Nα in the cyclobutyl glycine residue (**Figure 5A**, compound **36**). Since the incorporation of a methyl group at this position disrupts a key hydrogen bond with Leu49 in CBX- CDs (Stuckey et al., 2016a), as expected, **36** showed no cellular activity in the CBX8 reporter assay (**Figure 5B**). However, once again, **36** was more toxic than UNC7040 with a DC_50_ of 3.6 ± 1.8 µM. As an alternative approach, we incorporated an Nα-methylated valine at the alanine position (**37**), but **37** was again more toxic than UNC7040 (**Figure 5C**, DC50 = 4.4 ± 0.41 µM) but inactive as expected in the CBX8 reporter assay (**Figure 5B**). Finally, we focused our search for a nontoxic negative control on the lysine mimetic position. As depicted in **Figure 4**, close structural analogues, **19** and **22**, that differ only in bearing a methyl versus isobutyl group, respectively, differ dramatically in their cellular activity (**22**, EC50 = 2.8 ± 0.49 µM whereas **19**, EC50 >100 µM). Based on this result, we synthesized compound **38** (UNC7263), which is the methyl, norbornyl lysine mimetic analogue of UNC7040, which possesses no cellular activity in the CBX8 reporter assay as well as no cytotoxicity up to 100 µM (**Figure 5A**, **B, C**). Thus, even though the *in vitro* potency for CBX8 in our TR-FRET competition assay is identical for UNC7040 versus UNC7263 (**Figure 6A**), and their cell permeability is expected to be identical (**Figure S2H,** compare **30** and **33**), the PAM activity of UNC7040 results in potent cellular activity that is attenuated by at least 100-fold in UNC7263. While creation of a negative control through modulation of allostery in a ternary complex is atypical, it is perhaps a no less valid comparator than negative controls that lack affinity for the target protein of interest. The fact that our attempts to produce loss of affinity negative controls led to toxic compounds highlights the chief logical disconnect for all negative controls; a substituent that blocks binding to the target of interest may introduce new activity not present in the positive control. The mechanism underlying the toxicity of **35**, **36**, and **37** is unknown, and since it cannot be attributed to CBX-CD binding, it was not investigated further. With final probe, UNC7040 and negative control, UNC7263 in hand, we proceeded with more thorough profiling of selectivity and biological activity.

**Figure 6.**
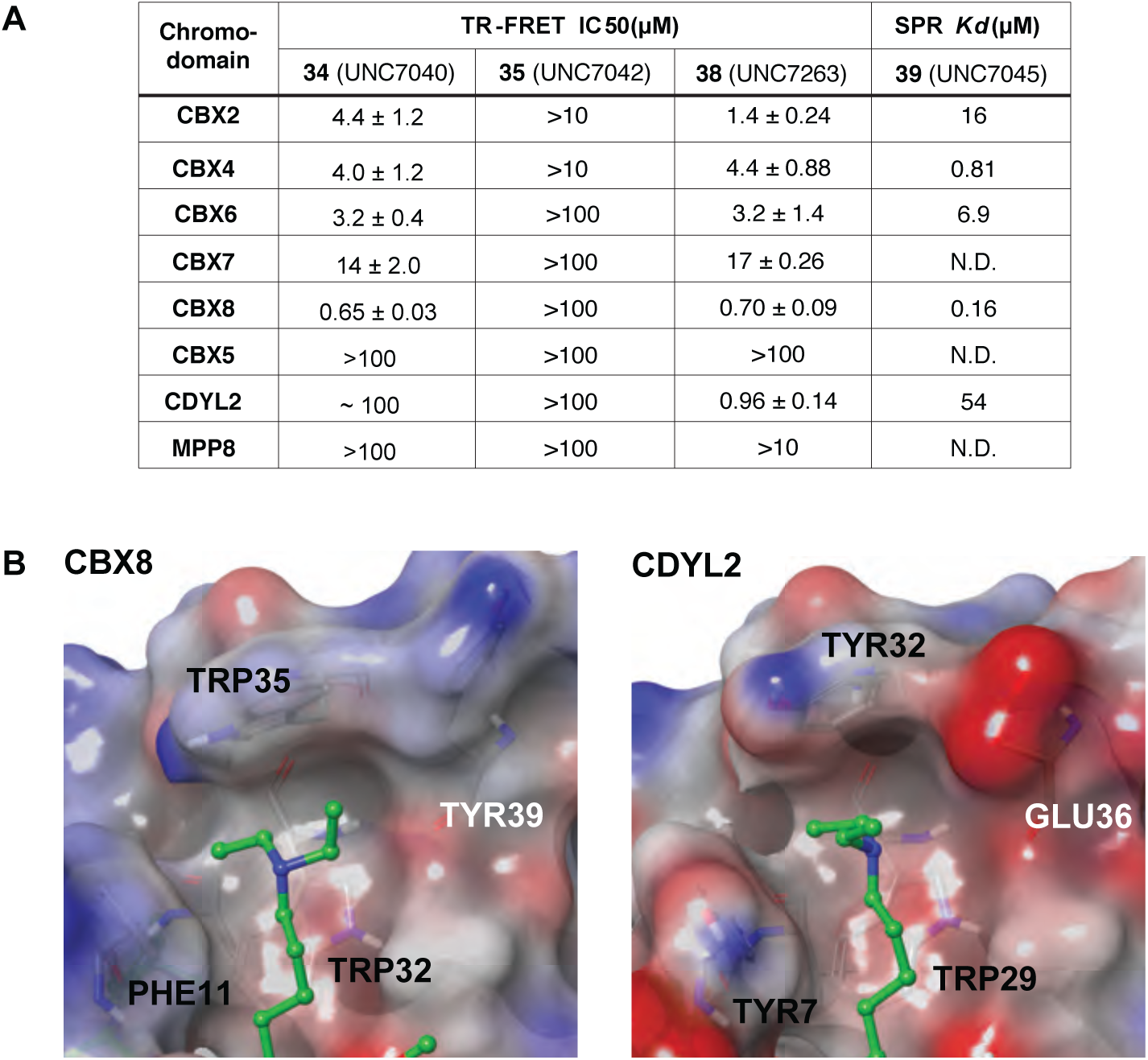
Quantitative analysis of 34, 35, 38 and 39 binding to CBXs, CDYL2, and MPP8 chromodomains by TR-FRET and SPR and structural comparison of the aromatic cages of CBX8 and CDYL2. (A) The binding profile of 34 (UNC7040), 35 (UNC7042), 38 (UNC7263) and 39 (UNC7045) for CBXs, CDYL2, and MPP8 chromodomains. For CBX2 and CBX4, top two concentrations were discarded for technical issues in TR-FRET assay. Data shown are mean ± SD, n = 6. (B) The side chain of Tyr39 in CBX8 faces out from the aromatic cage making it more expansive than that of CDYL2 (right). The side chain of Glu36 in CYDL2 facing into the aromatic cage makes it more compact (left).

### In Vitro Selectivity Profiling of UNC7040

We have previously used protein domain microarrays to evaluate the specificity of chemical probes that target methyllysine-binding domains like MBTs, CDs and Tudors (Bae et al., 2017; James et al., 2013; Lamb et al., 2019; Stuckey et al., 2016a). We evaluated the binding specificity of our CBX8 PAM UNC7040, in a similarly broad and unbiased fashion versus a methyl- lysine binding domain array containing 274 purified Kme readers, including 33 CDs, 43 Tudor domains, >100 PHDs as well as representatives from a number of additional domains such as PWWP, BAH, ELM2, HORMA domains, and Ank repeats. This protein microarray contains GST fusions of these domains immobilized on a glass slide (see, **S5B** for list of the domains). Biotin linked analogue of UNC7040, compound **39** was then pre-conjugated to streptavidin-Cy3 and used to probe these arrays. In this assay, the tagged probe bound strongly to CBX2 and CBX8 CDs, and also interacted less strongly with CBX4, CBX6 and CBX7, and weakly to the CD of CDYL2 (**Figure S5A**). As expected, **39** did not display binding to CBX1, CBX3, or CBX5 which are CBX CDs of human heterochromatin protein 1 (HP1) paralogs. This qualitative data focused our quantitative selectivity profiling on CBX domains.

To quantitate the selectivity of UNC7040 we utilized a TR-FRET assay with CBX2/4/6/5/7/8, CDYL2, and MPP8 CD which was recently established (Rectenwald et al., 2019). The IC50 trends generated from TR-FRET results were generally consistent with the intensity of the microarray data (**Figure 6A**, UNC7040). UNC7040 is more than ∼5-fold selective for CBX8 against CBX2, CBX4 and CBX6, and 22-fold selective for CBX8 over CBX7 via TR-FRET analysis. We also tested compound **35**, which represents an *in vitro* negative control for UNC7040, and UNC7263, which is the cellular negative control for UNC7040. As expected, **35** did not bind to the CDs in this study at concentrations up to 100 µM (**Figure 6A**, **35**) and UNC7263, which still has the potential to show *in vitro* binding to these proteins, displayed a similar affinity profile to UNC7040 (**Figure 6A**, UNC7263). The only significant difference between UNC7040 and UNC7263 is the binding affinity to CDYL2, with IC50’s of ∼ 100 µM and 0.96 ± 0.14 µM, respectively. Based on the UNC3866-bound crystal structures of CBX8 and CDYL2, the aromatic cage of CDYL2 is more constrained compared to that of CBX8 because of the side chain of Glu36 in CDYL2 facing into the aromatic cage while the side chain of Tyr29 in CBX8 faces out from the aromatic cage (**Figure 6B**). Therefore, UNC7263 would more favorably bind to CDYL2 compared to UNC7040 which has a bulkier side chain at the lysine mimetic position.

In addition to the TR-FRET assay, we utilized surface plasmon resonance (SPR) to measure *Kd* values of compound UNC7040 versus different CDs including CBX2/4/6/8 and CDYL2. To achieve this, we used compound **39,** a biotinylated derivative of UNC7040, and immobilized this compound on a NeutrAvidin chip. The trends in Kd generated from SPR were consistent with the TR-FRET data except CBX2 which showed a little higher *Kd* (**Figure 6A**). Compound **39** is more than 5-fold selective for CBX8 against CBX4, 43-fold selective for CBX8 over CBX6, and more than 100-fold selective over CBX2 and CDYL2. Taken together, *in vitro* binding data suggests that compound UNC7040 possessed at least 5-fold *in vitro* selectivity within CBX CDs and excellent selectivity against other methyl-lysine binding domains. Given the high homology of PRC1 CBX domains, the selectivity achieved within this family is perhaps as good as can be expected.

### Cell Reporter Selectivity Profiling of UNC7040

We next sought to assess the selectivity of compound UNC7040 against another CD in a cellular context by utilizing a previously reported CBX7 GFP reporter assay (Lamb CCB 2019). Here, as a positive control, UNC4976 was tested along with compound UNC7040, and consistent with our published data, UNC4976 had an EC50 of 3.0 ± 0.40 µM (n = 9). On the contrary, UNC7040 demonstrated no cellular activity in the CBX7 reporter cell line (**Figure S5C**).

### Activity of UNC7040 in clinically relevant cancer cell lines

To assess the activity of UNC7040 in a CBX8-dependent, clinically relevant cancer model, we interrogated the impact of treatment on two different cancer types; diffuse large B-cell lymphoma (DLBCL) and colorectal cancer (CRC). Lymphomagenesis results from aberrant transcriptional repression of cell-cycle check point and B-cell maturation genes by the polycomb pathway (Béguelin et al., 2016). Specifically, silencing is mediated by PRC1 containing BCOR and CBX8. By binding H3K27me3, CBX8 stabilizes BCOR interaction with BCL6 and facilitates robust transcriptional silencing (Béguelin et al., 2016). Importantly, CBX8 RNAi knockdown mimics the antiproliferative effects of EZH2 and BCL6 inhibition in DLBCL cells arguing that CBX8 presents a potential therapeutic target in B-cell lymphoma. To evaluate the impact of small molecule-dependent CBX8 antagonism, we examined proliferation of SUDHL4 cells in response to UNC7040 or UNC7263 treatment. To benchmark against proliferation defects triggered by polycomb inactivation, we also treated SUDHL4 cells with small molecules targeting PRC2, UNC1999 (EZH2 inhibitor) or EED226 (EED antagonist). As expected, small molecule-mediated PRC2 inactivation, but not the respective control compounds (UNC2400, UNC5679), dramatically reduced DLBCL cell proliferation (**Figure 7A**). Importantly, SUDHL4 cell proliferation was also impaired when treated with UNC7040, but not UNC7263, consistent with CBX8 acting downstream of PRC2. In addition, we observed that cell proliferation was further reduced by combined treatment of UNC7040 with EED226 suggesting that allosteric PRC1 and PRC2 inhibition have additive effects on DLBCL cell growth (**Figure 7B**).

**Figure 7.**
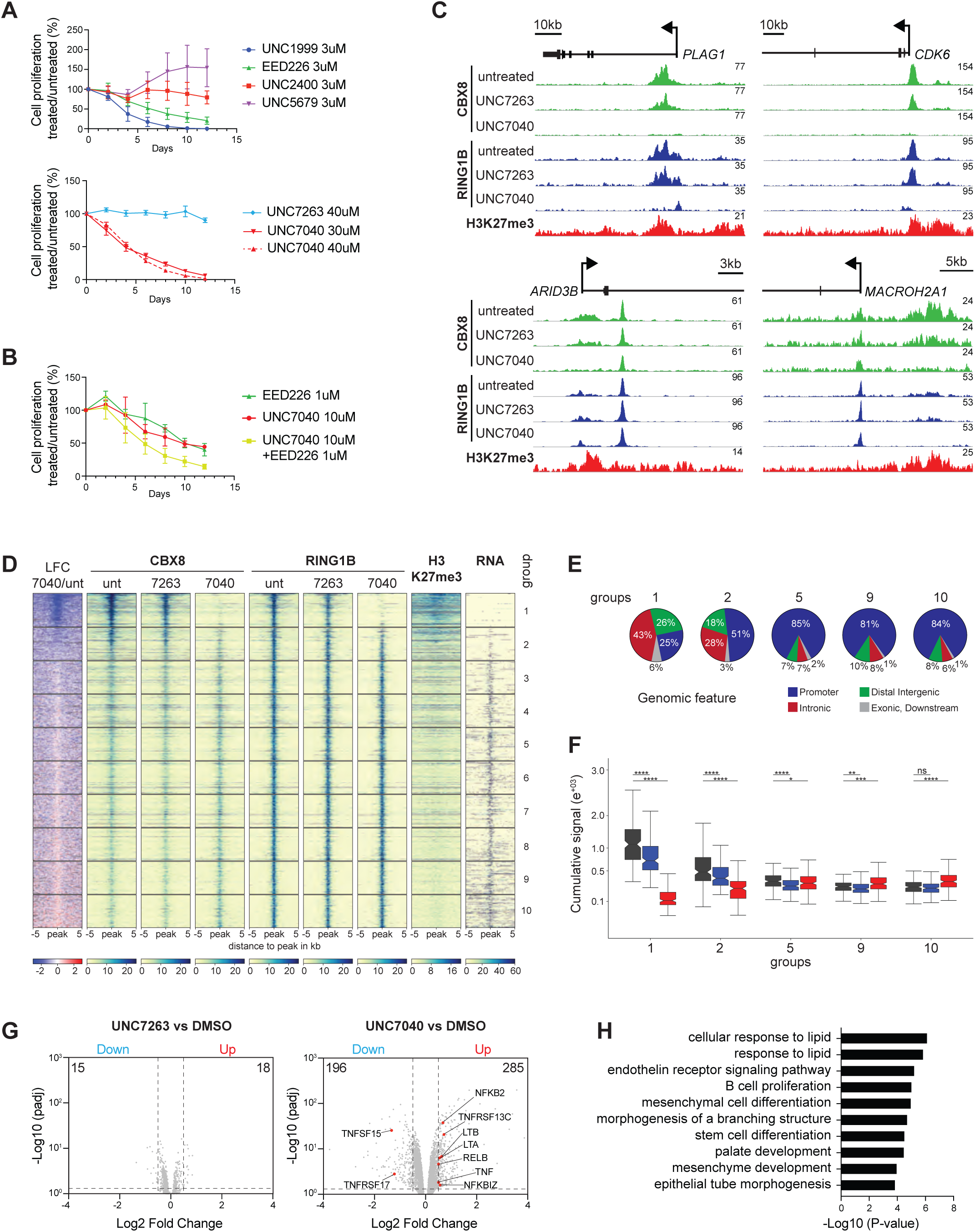
UNC7040 causes reduced DLBCL cell proliferation and activation of B-cell maturation genes by blocking canonical PRC1 targeting via CBX8. (A) Effects of treatment with PRC2 (UNC1999, EED226) or CBX8 (UNC7040) antagonists on SUDHL4 cell proliferation relative to vehicle control. UNC2400, UNC5679 and UNC7263 served as negative controls for UNC1999, EED226 and UNC7040, respectively. (B) Comparison of individual and combined treatment with UNC7040 and EED226 on SUDHL4 cell proliferation. Data in (A) and (B) represents an average of four to six replicates ± standard deviation. (C) Genomic screenshots of CBX8 and RING1B ChIPseq signals at four selected PRC2 target genes in SUDHL4 cells treated for six hours with DMSO, UNC7263 or UNC7040. Bottom track shows H3K27me3 signal in untreated cells. (D) Heatmaps display ChIPseq enrichment of CBX8 and RING1B in SUDHL4 cells treated for 6 h with DMSO, UNC7263 at 40 μM or UNC7040 at 40 μM. Also shown are signals of H3K27me3 and mRNA in untreated cells. The signals from two independent ChIPseq experiments are centered around RING1B peaks ± 5 kb and plotted as spike-in normalized mapped read counts. Genomic intervals were separated into 10 equal groups and sorted based on log2 fold change in CBX8 occupancy upon UNC7040 treatment relative to DMSO. (E) Pie charts display relative distribution of genomic features in selected cChIP-seq groups based on (D). (F) Box plots compare cumulative CBX8 signal (±0.5 kb around RING1B peaks) for selected groups in untreated and treated SUDHL4 cells. Significance (P value) was calculated using Wilcoxon signed-rank test (*: p <= 0.05; **: p <= 0.01; ***: p <= 0.001; ****: p <= 0.0001; ns, not significant). (G) Volcano plots show gene expression changes of UNC7263-treated (left) or UNC7040-treated SUDHL4 cells relative to DMSO. Numbers in represent repressed (blue) and upregulated (red) genes. Differential gene expression (P_adj_ < 0.05; LFC: <-0.5 and >0.5, respectively) was calculated from three independent replicates. Highlighted are genes associated with B-cell maturation. (H) For functional analysis of canonical CBX8-PRC1 targets, differential peaks resulting from UNC7040 treatment were analyzed by GREAT (v.4.0.4) using the nearest gene within 100 kb to generate enrichment of biological processes.

To relate the proliferation defect in response to UNC7040 to changes in PRC1 occupancy, we used calibrated Chromatin Immunoprecipitation coupled to next-generation sequencing (cChIP-seq). Compound treatment was limited to six hours to exclude indirect effects on PRC1 binding as a consequence of transcriptional changes. In addition, we profiled gene expression changes by RNA-seq after six days. At first, we used cChIP-seq of CBX8, RING1B and H3K27me3 in untreated cells to determine the genome-wide distribution of PRC1 and PRC2 targets. We detected 1735 peaks with significant RING1B enrichment facilitating identification of cPRC1 and vPRC1 binding sites in SUDHL4 cells (**Figure 7C, D**). Similar to other cell types (Fursova et al., 2019; Scelfo et al., 2019; Zepeda-Martinez et al., 2020), only a fraction of RING1B overlapped with H3K27me3 suggesting that most PRC1 targeting involves PRC2-independent mechanisms (groups 3-10, vPRC1 targets). CBX8 was preferentially enriched at H3K27me3 targets, consistent with its proposed role in cPRC1 recruitment (groups 1 and 2, cPRC1 targets) (**Figure 7D**). Unlike in mESCs, H3K27me3 peaks were located predominantly at intronic and distal intergenic regions suggesting that PRC2 signals for CBX8 recruitment at enhancers in DLBCL cells (**Figure 7E)**. Short treatment with UNC7040 strongly reduced CBX8 and RING1B occupancy at H3K27me3-modified regions (groups 1 and 2), whereas binding remained largely unaffected at most vPRC1 target sites (groups 3-8) (**Figure 7C, D, F**). Surprisingly, at vPRC1 targets in groups 9 and 10, CBX8 binding was modestly but significantly increased upon UNC7040 treatment (**Figure 7D and F**). By comparison, UNC7263 caused less cPRC1 displacement and only marginal changes in CBX8 occupancy at vPRC1 targets, consistent with its limited effect on SUDHL4 cell proliferation and lack of activity in our CBX8 reporter assay (**Figures 5B, 7A**). These results demonstrate that UNC7040 more efficiently disrupts H3K27me3 recognition and canonical CBX8 function in cancer cells, similar to reporter mESCs. Based on increased CBX8 binding at vPRC1 targets, UNC7040 may reinforce cPRC1 eviction *in vivo* by promoting CBX8 recruitment to non-canonical target loci. It remains unknown whether genomic CBX8 redistribution is only linked to PAM activity enhancing nucleic acid binding.

Transcriptome analysis by RNA-seq revealed that UNC7040 triggered robust, differential expression of 481 genes (padj. 0.05, LFC=+/-0.5; **Figure 7G and Figure S6A**). Most genes with altered mRNA levels were upregulated in line with loss of polycomb repression by UNC7040. Given its limited impact on cPRC1 occupancy, UNC7263 treatment had negligible impact on SUDHL4 gene regulation. Importantly, UNC7040-induced genes included members of the Tumor necrosis factor (TNF) superfamily and regulators involved in NF-κB signaling (**Figure 7G**). Both pathways are important for promoting B cell maturation (Mackay and Schneider, 2009), consistent with reduced proliferation of SUDHL4 cells upon UNC7040 treatment. Being enriched at distal intergenic and intronic regions marked by H3K27me3, CBX8-PRC1 might control B-cell maturation genes by repressing the activity of regulatory enhancers. To predict genes under control of CBX8-dependent enhancers, we employed Genomic Regions Enrichment of Annotations Tool (GREAT (McLean et al., 2010)). GREAT analysis of UNC7040-sensitive CBX8 peaks revealed enrichment of genes with functional annotation in B-cell proliferation supporting the notion that CBX8 directs cPRC1-dependent silencing to enhancers of B-cell maturation genes and this control is disrupted by UNC7040 (**Figure 7H**).

CRC present another emerging model for CBX8-dependent control of cell proliferation, but the underlying mechanism remains largely unexplored (Zhang et al., 2019). We selected a panel of established CRC cell lines, confirmed CBX8 expression and evaluated changes in proliferation after 72 hours of treatment with either vehicle control or with increasing concentrations of UNC7040 or UNC7263 (**Figure S6B, C**). UNC7040 treatment reduced proliferation of LoVo cells at concentrations above 1.85 μM, indicating CBX8 dependency of this CRC cell line (**Figure S6C**). The antiproliferative effects were specific to CBX8 inhibition since treatment with CBX7 antagonist UNC4976 or UNC7263 displayed cell counts similar to vehicle control (**Figure S6C, D**). UNC7040 treatment also impaired spheroid formation of LoVo cells but not HCT116 cells, corroborating the cell line-specific effect (**Figure S6E**).

Similar to DLBCL cells, cChIP-seq profiling revealed strong reduction in CBX8 and RING1B occupancy at H3K27me3 targets in LoVo cells, indicating efficient cPRC1 displacement in response to UNC7040 (**Figure S6F, G**). Despite substantial cPRC1 loss, UNC7040 triggered only minor transcriptional changes suggesting that additional mechanisms such as vPRC1 might act redundantly to maintain gene silencing in CRC cells (**Figure S6H, I**). Nevertheless, we found that expression of *KRT20*, a differentiation marker gene, was upregulated whereas LGR5, a stem cell marker gene, was downregulated by UNC7040, consistent with CBX8-dependence of LoVo cell proliferation and self-renewal (Shimokawa et al., 2017). In conclusion, epigenomic and transcriptomic profiling in two distinct cancer cell lines demonstrated potent activity of UNC7040 to displace CBX8-containing cPRC1 and impair polycomb-dependent gene silencing.

## CONCLUSIONS

Here, we took advantage of a synthetic cellular reporter assay of CBX8-dependent repression and an iterative, structure-guided approach to design UNC7040, and a negative control, UNC7263. Strikingly, while these compounds have similar *in vitro* affinity for CBX8, their cellular activity differs dramatically. This difference can be explained by UNC7040’s enhanced ability to stabilize non-specific binding of CBX8 to nucleic acids as a PAM. Alternatively, the ability of UNC7040 to engender unique CBX8 conformations, as observed by NMR and MD, may influence interactions within PRC1 or non-canonical complexes in ways that can’t be recapitulated *in vitro*. This possibility will be the subject of future studies.

Application of UNC7040 in two distinct cancer types demonstrated rapid and efficient displacement of CBX8-containing PRC1 from facultative heterochromatin marked with H3K27me3. In contrast, CBX8 and RING1B remained largely unchanged at non-canonical targets. Notably, in DLBCL cells CBX8 was modestly increased at a fraction of vPRC1 targets, consistent with enhanced interactions of CBX8 with vPRC1 subunits upon UNC7040 treatment. It remains to be tested if alternative PRC1 interactions are coupled to PAM-induced nucleic acid binding. In any case, these results support previous observations of CBX8 engaging with diverse polycomb-dependent and independent chromatin-modifying complexes (Béguelin et al., 2016; Chung et al., 2016; Creppe et al., 2014; Morey et al., 2012). Moreover, many of its non-canonical targets are active promoters suggesting that in addition to transcriptional repression, CBX8 may play a role in promoting gene expression that remains to be explored.

By blocking interaction with H3K27me3, UNC7040 offers a powerful new molecular tool to investigate the distinct functions of CBX8 in polycomb-dependent and independent gene regulation. The combinatorial nature of PRC1 assembly and its partial redundancy have hampered functional dissection of canonical and variant PRC1 subunits using classical genetic approaches. Adding to a chemical biology tool kit including small molecule-induced degraders of polycomb group proteins (Dobrinić et al., 2020; Hsu et al., 2020; Ma et al., 2020; Potjewyd et al., 2020; Zepeda-Martinez et al., 2020), UNC7040 provides an opportunity for selective, acute and reversible inactivation of CBX8 reader activity to distinguish its role in canonical PRC1 regulation. The versatility of UNC7040 to examine canonical CBX8 function without laborious genetic manipulations will be very impactful to study its functions in gene regulation of different cell types and tissues, in normal development and disease. Furthermore, a CBX8-specific PAM chemical probe is highly relevant for translational studies to explore the potential of CBX8 as target for cancer therapy. Within the framework of the Structural Genomics Consortium chemical probes effort (Müller et al., 2018), we will make UNC7040 available to the expert biomedical community for use in *in vitro* and cellular disease models.

## AUTHOR CONTRIBUTIONS

Conceptualization, J.L.S., L.I.J., S.V.F. and O.B.; Formal Analysis; D.B., D.B.K., T.M.W. and J.L.S.; Investigation, J. L. S., D.B., Y.S., T.M.W., B.H., C.P., R.L., S.S., H.O., J.M.R., J.L.N., S.H.C., C.S., J.D.U. and B.H.; Writing – Original Draft, J.L.S., D.B., T.M.W., C.A.M., D.B.K., O.B. and S.V.F.; Writing – Review & Editing, J.L.S., D.B., B.H., M.T.B., H.J.L, S.M.M., J.M.K., G.G.W, K.H.P., D.B.K., L.I.J., C.A.M., S.V.F. and O.B.; Resources, M.T.B., H.J.L, S.M.M., J.M.K., G.G.W, K.H.P., D.B.K., L.I.J., C.A.M., S.V.F. and O.B.; Visualization, J.L.S., D.B., B.H., T.M.W., D.B.K. and O.B.; Supervision, M.T.B., H.J.L, S.M.M., J.M.K., G.G.W, K.H.P., D.B.K., L.I.J., C.A.M., S.V.F. and O.B.; Project Administration, O.B. and S.V.F.; Funding Acquisition, L.I.J., O.B., M.T.B. and S.V.F.

## ACKNOWLEDGMENTS

This work was supported by the by National Cancer Institute, National Institutes of Health (NIH) (R01CA218392) to S.V.F. and O.B.; National Institute of General Medical Sciences, NIH (R01GM100919) to S.V.F., (R35GM128705) to C.A.M. and (R01GM132299) to D.B.K.; by the Austrian Academy of Sciences, the New Frontiers Group of the Austrian Academy of Sciences (NFG-05) and Norris Comprehensive Cancer Center, Keck School of Medicine, USC to O.B.; by National Institute on Drug Abuse NIH (R61DA047023) to L.I.J.; NCI P30CA014089, Gloria Borges WunderGlo Foundation, Gene Gregg Pancreas Cancer Research Fund to H.J.L.; by The Holden Comprehensive Cancer Center at The University of Iowa and its National Cancer Institute Award P30CA086862 to C.A.M.; by the UNC Lineberger Comprehensive Cancer Center and its National Cancer Institute Award P30CA016086 to S.V.F; and by the UNC Eshelman Institute for Innovation (RX0351210 and RX03712105) to D.B.K. Probing of arrayed methyl-binding domains was made possible via the UT MDACC Protein Array & Analysis Core (PAAC) CPRIT (RP180804) to M.T.B. We acknowledge the UNC Longleaf supercomputer cluster and their staff for support. The authors thank Joshua Kritzer (Tufts) for sharing the CAPA HeLa cell line, and Isabelle Engelberg, Catherine Foley and Frances Potjewyd for reviewing primary data supporting this manuscript.

## EXPERIMENTAL MODEL AND SUBJECT DETAILS

### Cell Lines

#### Generation of Polycomb in-vivo Assay in mouse Embryonic Stem Cell (mESC) and Culture Conditions

CBX8 reporter mESCs with 12xZFHD1, 4xGAL4 and 7xTETO DNA binding sites upstream of a Puromycin-GFP reporter gene were generated previously described dual reporter mESCs (Moussa et al., 2019) by recombinase mediated cassette exchange. mESCs were cultivated without feeders in high-glucose-DMEM (Sigma, D6429) supplemented with 13.5% fetal bovine serum (Gibco), 10 mM HEPES pH 7.4 (Corning, 25-060-CI), 2 mM GlutaMAX (Gibco, 35050- 061), 1 mM Sodium Pyruvate (Gibco, 11360-070), 1% Penicillin/Streptomycin (Sigma, P0781), 1X non-essential amino acids (Gibco, 11140-050), 50 mM β-mercaptoethanol (Gibco, 21985-023) and recombinant LIF, and incubated at 37°C and 5% CO2. mESCs were passaged every 48 hours by trypsinization in 0.25% Trypsin-EDTA (1X) (Gibco, 25200-056) and seeding of 2.0 x 10^6^ cells on a 10 cm tissue culture plate (Genesee Scientific, #25-202).

### Culturing of cancer cell lines

Human DLBCL cells, SUDHL4 (ATCC #CRL-2957), were cultured in RPMI-1640 complemented with 10% fetal bovine serum (Gibco) and 1% penicillin/streptomycin. The human colorectal cell lines HCT116 and HT-29 (ATCC #CCL-247 and #HTB-38, respectively) were cultured in McCoy’s 5A medium supplemented with 10% fetal bovine serum (Gemini) and 1% penicillin/streptomycin (Gemini). Caco2 (ATCC #HTB-37) were maintained in EMEM supplemented with 10% fetal bovine serum and 1% penicillin/streptomycin. LoVo (ATCC #CCL-229) were maintained in F-12K medium supplemented with 10% fetal bovine serum and 1% penicillin/streptomycin. The DLD-1 cell line (gift from Dr. Yun at Baylor College of Medicine) was cultured in McCoy’s 5A medium supplemented with 10% fetal bovine serum and 1% penicillin/streptomycin.

## METHOD DETAILS

### Protein Expression and Purification

#### Expression constructs

The chromodomains of CBX2 (residues 9–66 of NP_005180), CBX4 (residues 8–65 of NP_003646), CBX6 (residues 8–65 of NP_055107), CBX7 (residues 8–62 of NP_783640) and CDYL2 (residues 1–75 of NP_689555) were expressed with C-terminal His-tags in pET30 expression vectors. The chromodomain of CBX5 (residues 18–75 of NP_036429) and MPP8 (residues 55–116 of NP_059990) were expressed with a N-terminal His-tag in a pET28 expression vector. The chromodomain of CBX8 (residues 8–61 of NP_065700) was expressed with either a N-terminal His-tag in a pET28 expression vector or a N-terminal GST-tag in a pGEX derived expression vector.

#### Protein expression and purification

All expression constructs were transformed into Rosetta BL21(DE3)pLysS competent cells (Novagen, EMD Chemicals, San Diego, CA). Protein expression was induced by growing cells at 37 °C with shaking until the OD600 reached ∼0.6–0.8 at which time the temperature was lowered to 18 °C and expression was induced by adding 0.5 mM IPTG and continuing shaking overnight. Cells were harvested by centrifugation and pellets were stored at −80 °C.

His-tagged proteins were purified by re-suspending thawed cell pellets in 30 ml of lysis buffer (50 mM sodium phosphate pH 7.2, 50 mM NaCl, 30 mM imidazole, 1× EDTA free protease inhibitor cocktail (Roche Diagnostics, Indianapolis, IN)) per liter of culture. Cells were lysed on ice by sonication with a Branson Digital 450 Sonifier (Branson Ultrasonics, Danbury, CT) at 40% amplitude for 12 cycles with each cycle consisting of a 20 s pulse followed by a 40 s rest. The cell lysate was clarified by centrifugation and loaded onto a HisTrap FF column (GE Healthcare, Piscataway, NJ) that had been pre-equilibrated with 10 column volumes of binding buffer (50 mM sodium phosphate, pH 7.2, 500 mM NaCl, 30mM imidazole) using an AKTA FPLC (GE Healthcare, Piscataway, NJ). The column was washed with 15 column volumes of binding buffer and protein was eluted in a linear gradient to 100% elution buffer (50 mM sodium phosphate, pH 7.2, 500 mM NaCl, 500 mM imidazole) over 20 column volumes. Peak fractions containing the desired protein were pooled and concentrated to 2 ml in Amicon Ultra-15 concentrators 3,000 molecular weight cut-off (Merck Millipore, Carrigtwohill Co. Cork IRL). Concentrated protein was loaded onto a HiLoad 26/60 Superdex 75 prep grade column (GE Healthcare, Piscataway, NJ) that had been pre-equilibrated with 1.2 column volumes of sizing buffer (25 mM Tris, pH 7.5, 250 mM NaCl, 2 mM DTT, 5% glycerol) using an ATKA Purifier (GE Healthcare, Piscataway, NJ). Protein was eluted isocratically in sizing buffer over 1.3 column volumes at a flow rate of 2 ml/min collecting 3-ml fractions. Peak fractions were analyzed for purity by SDS-PAGE and those containing pure protein were pooled and concentrated using Amicon Ultra-15 concentrators 3,000 molecular weight cut-off (Merck Millipore, Carrigtwohill Co. Cork IRL). Protein was exchanged into a buffer containing 25 mM Tris, pH 7.5, 150 mM NaCl, 2 mM β-mercaptoethanol before use in ITC.

GST-tagged CBX8 was purified by re-suspending thawed cell pellets in 30 ml of lysis buffer (1× PBS, 5 mM DTT, 1× EDTA free protease inhibitor cocktail (Roche Diagnostics, Indianapolis, IN)) per liter of culture. Cells were lysed on ice by sonication as described for His-tagged proteins. Clarified cell lysate was loaded onto a GSTrap FF column (GE Healthcare, Piscataway, NJ) that had been pre-equilibrated with 10 column volumes of binding buffer (1x PBS, 5mM DTT) using a AKTA FPLC (GE Healthcare, Piscataway, NJ). The column was washed with 10 column volumes of binding buffer and protein was eluted in 100% elution buffer (50 mM Tris, pH 7.5, 150 mM NaCl, 10 mM reduced glutathione) over 10 column volumes. Peak fractions containing the desired protein were pooled and concentrated to 2 ml in Amicon Ultra-15 concentrators, 10,000 molecular weight cut-off (Merck Millipore, Carrigtwohill Co. Cork IRL). Concentrated protein was loaded onto a HiLoad 26/60 Superdex 200 prep grade column (GE Healthcare, Piscataway, NJ) that had been pre-equilibrated with 1.2 column volumes of sizing buffer (25 mM Tris, pH 7.5, 500 mM NaCl, 2 mM DTT, 5% glycerol) using an ATKA FPLC (GE Healthcare, Piscataway, NJ). Protein was eluted isocratically in sizing buffer over 1.3 column volumes at a flow rate of 2 ml/min collecting 3 mL fractions. Peak fractions were analyzed for purity by SDS-PAGE and those containing pure protein were pooled and concentrated using Amicon Ultra-15 concentrators 10,000 molecular weight cut-off (Merck Millipore, Carrigtwohill Co. Cork IRL).

#### Affinity tag removal

The N-terminal GST-tag was removed from CBX8 proteins by TEV protease cleavage according to manufacturer’s recommendations (Sigma-Aldrich, St. Louis, MO). Briefly, purified protein was incubated with TEV protease at a final concentration of 50 units TEV protease per milligram tagged protein for 16 h at 4 °C. The cleavage reaction was loaded onto a HiLoad 26/60 Superdex 75 to separate tag free CBX8 from GST and any protein that still retained the GST-tag. Size exclusion was performed as described above. Peak fractions were analyzed for purity by SDS-PAGE and those containing pure tag free CBX8 protein were pooled and concentrated using Amicon Ultra-15 concentrators 3,000 molecular weight cut-off (Merck Millipore, Carrigtwohill Co. Cork IRL). Protein was exchanged into a buffer consisting of 20mM MES pH 6.5, 250mM NaCl, 1mM DTT.

### TR-FRET Assay

TR-FRET assay was performed as described in Rectenwald *et al*. (Rectenwald et al., 2019). A stock solution of 10X Kme reader buffer (200 mM Tris pH 7.5, 1500 mM NaCl, and 0.5% Tween 20) was prepared, 0.2 µm filtered, stored at room temperature, and was used throughout. Assays were completed using freshly made Kme reader buffer containing 20 mM Tris pH 7.5, 150 mM NaCl, 0.05% Tween 20, and 2 mM dithiothreitol (DTT). White, low-volume, flat-bottom, non-binding, 384-well microplates (Greiner, #784904) were used for assay development and screening with a total assay volume of 10 µL. 384-well, V-bottom polypropylene plates (Greiner, #781280) were used for compound serial dilutions and for transfer of assay mixtures. For compounds with stock solutions in water, serial dilutions were made using Kme reader buffer. For compounds stored in DMSO, serial dilutions were made using DMSO. Following addition of all assay components, plates were sealed with clear covers, gently mixed on a tabletop shaker for 1 minute, centrifuged at 1000 x g for 2 minutes, and allowed to equilibrate in a dark space for one hour before reading. Measurements were taken on an EnVision® 2103 Multilabel Plate Reader (Perkin Elmer) using an excitation filter at 320 nm and emission filters at 615 nm and 665 nm. 615 nm and 650 nm emission signals were measured simultaneously using a dual mirror at D400/D630. TR-FRET output signal was expressed as emission ratios of acceptor/donor (665/615 nm) counts. Percent inhibition was calculated on a scale of 0% (i.e., activity with DMSO vehicle only) to 100% (100 µM UNC3866) using full column controls on each plate. The interquartile mean of control wells was used to calculate Z’ values. For dose-response curves, data was fit with a four-parameter nonlinear regression analysis using GraphPad Prism 7.0 or ScreenAble software to obtain IC_50_ values.

For testing compounds in higher throughput, 384-well assay ready plates were prepared in standard plate format: columns 1 and 2 were used for low signal controls (100% inhibition with competitor compound), columns 23 and 24 were used for high signal controls (DMSO only), and columns 3–22 were used for 25 µM single-dose test compounds. First, controls were added to a mother plate where columns 1 and 2 were filled with 10 mM stock of UNC3866 in DMSO and columns 23 and 24 were filled with DMSO. Test compounds were dispensed across the mother plate at 100X (10 mM) concentration in columns 3–22 using a TECAN Freedom EVO liquid handling workstation. Using a TTP Labtech Mosquito® HTS liquid handling instrument, assay ready plates were stamped by stamping 100 nL of control compound into columns 1 and 2, 25 nL of compounds from the mother plate into columns 3–22, and 25 nL of DMSO into columns 23 and 24. Protein, biotinylated tracer ligand, and the TR-FRET reagents were added together and gently mixed by pipetting and rocking. 10 µL was then added to each well of an assay ready plate using a Multidrop Combi (ThermoFisher). Percent inhibition was calculated on a scale of 0% (i.e., activity with DMSO vehicle only) to 100% (100 µM UNC3866) from the full column controls on each plate.

### CBX8-GFP Reporter Assay

CBX8-GFP reporter containing mESCs were cultivated without feeders in high-glucose-DMEM (Sigma, D6429) supplemented with 13.5% fetal bovine serum (Gibco), 10 mM HEPES pH 7.4 (Corning, 25-060-CI), 2 mM GlutaMAX (Gibco, 35050-061), 1 mM Sodium Pyruvate (Gibco, 11360-070), 1% Penicillin/Streptomycin (Sigma, P0781), 1X non-essential amino acids (Gibco, 11140-050), 50 mM β-mercaptoethanol (Gibco, 21985-023) and recombinant LIF, and incubated at 37°C and 5% CO_2_. mESCs were passaged every 48 hours by trypsinization in 0.25% Trypsin-EDTA (1X) (Gibco, 25200-056) and seeding of 2.0 x 10^6^ cells on a 10 cm tissue culture plate (Genesee Scientific, #25-202).

For the screening, compounds were prepared at 10 mM top concentration from 10 mM DMSO stocks as a 10-point and 2-fold dilution series. 0.5 mL of each serial-dilution was then added to a 384-well assay plate (Corning, 3764) in triplicate using a TTP Labtech Mosquito HTS liquid handling instrument. mESCs were trypsinized in 0.25% Trypsin-EDTA, counted on a BioRad TC20 cell counter, and diluted to a density of 2,000 cells/50 mL. 50 mL of cell suspension per well was then plated on top of previously added compound stocks to achieve a final 1X compound concentration + 0.99% DMSO, and assay plates were incubated for 48 hours at 37°C and 5% CO_2_. For testing the methyl ester version compounds, compounds were prepared at 10X concentration from 10 mM DMSO stocks as a 15-point, 1.5-fold dilution series, diluted into mESC media + 0.6% DMSO. 5 µL of each 10X stock was then added to a 384-well assay plate (Corning, 3764) in triplicate. mESCs were trypsinized in 0.25% Trypsin-EDTA, counted on a BioRad TC20 cell counter, and diluted to a density of 2,000 cells/45 µL. 45 µL of cell suspension per well was then plated on top of previously added 10X compound stocks to achieve a final 1X compound concentration + 0.06% DMSO, and assay plates were incubated for 48 hours at 37°C and 5% CO_2_.

After 48 hours, cells were washed once in 50 mL of 1X PBS and trypsinized with 12.5 mL of clear 0.25% Trypsin-EDTA (5X) (Gibco, 15400-054) per well. Cells were incubated at 37°C and 5% CO_2_ for 15–20 min to ensure in complete dislodging of cells from the assay plate. Trypsin was then quenched with 12.5 mL of 50% FBS in 1X PBS. Flow cytometry was completed on an Attune NxT equipped with Attune NxT v3.1 acquisition software. Live, single cells were gated for GFP expression and data analysis was completed with FlowJo and GraphPad Prism 8 software.

### Cell Toxicity Assay

The effect of DMSO and methyl ester version compounds (compound **34–38**) on cell viability was determined using a CellTiter-GloTM ATP detection system (Promega #7573). For testing DMSO tolerance, ten point, 1:0.8 and 1:0.5 dilution curves of DMSO starting 5 µM final concentration in mECS media were prepared. For testing compound tolerance, ten point, 1:2 dilution curves of compounds starting at 100 µM final concentration were diluted to 5X final concentration in mESC media. Then, 5 µL were added to 384-well white, clear bottom tissue culture plates (Corning #3707) with a Multimek automated liquid handling device (Nanoscreen, Charleston, SC). mESCs were trypsinized in 0.25% Trypsin-EDTA, counted on a BioRad TC20 cell counter, and diluted to a density of 5,000 cells/20 µL. 20 µL of cell suspension per well was then plated on top of previously added 5X DMSO or compound stocks to achieve a final 1X DMSO or compound concentration, and assay plates were incubated for 48 hours at 37°C and 5% CO_2_. After 48 hours, cells were lysed with 25 µL of CellTiter-GloTM reagent. Luminescence was read on an Envision platereader (Perkin Elmer) after 15 minutes at room temperature in dim light.

### EMSA Assay

All protein, DNA probe, and compound stocks were diluted to desired final concentrations in EMSA Binding Buffer (20 mM MES, pH 6.5, 250 mM NaCl, and 1 mM DTT). For protein titration experiments, a 2-fold serial dilution series was generated for CBX8 starting at 500 mM of protein. All protein samples were incubated with 500 nM DNA probe at a final reaction volume of 10 mL, on ice, for 20 minutes. For experiments with compounds, 15 mM of CBX8 was incubated with 500 nM DNA probe and a 2-fold serial dilution series of compounds starting at 25 µM of compound at a final reaction volume of 10 mL, on ice, for 20 minutes. All samples were diluted with 2 µL of 6X DNA loading dye (Thermo Scientific, #R0611) and loaded onto a Novex 10% TBE gels (EC62752BOX) and ran in 0.5X TBE buffer (pH 8.3, Novex TBE running buffer, #LC6675) at 100V and 4°C for 60 minutes, in the dark. Gels were imaged on a Multi-DocIt Imaging System with Doc- ItLS software, using a blue light plate to visualize FAM-labeled DNA probe fluorescence. After imaging, gels were then stained with SimplyBlue SafeStain (Invitrogen, LC6065) at room temperature for 2 hours.

### NMR Experiments

#### Protein expression and purification for NMR studies

The recombinant CBX8-CD was expressed in BL21 (DE3) (New England Biolabs) *Escherichia coli* cells. Cells were grown in LB-medium or M9-minimal media supplemented with ^15^N-NH_4_Cl or ^15^N-NH_4_Cl and ^13^C-glucose. For unlabeled protein, cells were grown shaking at 215 rpm at 37 °C until an OD ∼1.0 was reached and induced with 1 mM IPTG for 16–18 hr overnight. For isotopically-enriched protein, cells were grown in LB-medium until an OD ∼1.0, spun down at 4000 rpm for 10 minutes, and resuspended in M9-medium (4 l LB cells per 1 l M9) supplemented with either ^15^N-NH_4_Cl or ^15^N-NH_4_Cl/^13^C-glucose. The cells were allowed to recover in M9 media for 1 hr shaking at 18°C and induced with 1.0 mM IPTG for 16–18 hr overnight. Cells were subsequently collected by centrifugation at 6000 rpm for 20 minutes, frozen in N_2_(l) and stored at –80 °C.

For purification, cells were resuspended in a buffer containing 100 mM NaCl, 25 mM Tris (pH 7.5) supplemented with DNase I and lysozyme. Cells were then lysed using an Emulsiflex homogenizer (Avestin) or by sonication, and lysate cleared by centrifugation at 15 000 rpm for 1 hr at 4 °C. The soluble supernatant was incubated with glutathione agarose resin (ThermoFisher Scientific) rotating at 4 °C for 1 hr. The GST-tagged CBX8 CD was washed extensively with a high salt buffer containing 1 M NaCl and 25 mM Tris (pH 7.5) before elution with a buffer containing 150 mM NaCl, 25 mM Tris (pH 7.5) and 50 mM reduced glutathione. The GST-CBX8 CD was concentrated using a 3000 MWCO centrifugation filter unit to 2 ml and cleaved with TEV protease for 3 hr at 25 °C. The cleaved CBX8 CD was further purified using a combination of cation-exchange and size exclusion chromatography (Superdex 75, GE Healthcare Life Sciences). For NMR studies, ^15^N-CBX8 CD and ^15^N/^13^C-CBX8 CD were used in a final NMR buffer containing 100 mM NaCl and 40 mM phosphate buffer (pH 6.8). For EMSAs, the unlabeled CBX8 CD was used in a final buffer containing 25 mM phosphate buffer (pH 6.8), 25 mM NaCl, 1 mM EDTA, 1 mM DTT.

#### NMR experiments

Titrations experiments were performed by collecting ^15^N-HSQC spectra on 0.05 mM ^15^N-CBX8- CD in the apo state and upon addition of increasing concentrations of ligand. Titrations experiments were performed at 25 °C on a Bruker Avance II 800 MHz NMR spectrometer equipped with a 5 mm triple resonance cryoprobe. The data was processed using NMRPipe (Delaglio et al., 1995) and further analysis performed using CcpNmr (Vranken et al., 2005). To determine residues with CSPs, the following equation was used:

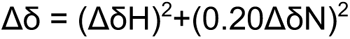

A resonance was considered significantly perturbed if the Δ*δ* value was greater than the average plus 0.5 standard deviations after trimming 10% of residues with the largest Δ*δ* value.

### CAPA Assay

HeLa cells stably expressing the HaloTag-GFP-Mito construct were provided by the Kritzer lab (Peraro et al., 2018; Peraro et al., 2017). Cells were cultured in high-glucose-DMEM (Sigma, D6429) supplemented with 10% fetal bovine serum, 1% Penicillin/Streptomycin (Sigma, P0781) and 1 mg/mL Puromycin (InvivoGen, ant-pr) and incubated at 37 °C and 5% CO_2_. Cells were passaged every 48–72 hours by trypsinization in 0.25% Trypsin-EDTA (1X) (Gibco, 25200-056) and seeding of 3.0 x 10^6^ cells on a T75 tissue culture plate.

GFP-HaloTag HeLa cells were seeded in a 384-well assay plate (Corning, 3764) at a density of 5,000 cells/well on the day before the experiment and allowed to adhere overnight at 37 °C and 5% CO_2_. A 100X mother plate of compound dilutions was prepared in a separate 384-well plate, and 10-points and 3-fold dilution series were generated using 10 mM DMSO stock. Compound-free DMSO control wells were also included in the mother plate to be used as no-pulse (100%) and no-pulse/no-chase (0%) controls. On the day of the experiment, a daughter plate of compound dilutions and control samples was prepared by transferring 0.5 µL of each well from the mother plate using a TTP Labtech Mosquito HTS liquid handling instrument. Then, the wells were diluted with 49.5 µL of HeLa media to ensure a final DMSO concentration of 1%. The media of the 384-well assay plate containing cells was aspirated and 50 μL of the compound and control solutions from the daughter plate were added to each well. The assay plate was then incubated at 37 °C and 5% CO_2_ for 4 hours. Media was removed, and cells were washed once with phenol-red free Opti-MEM (1X) (Gibco, 11058-021) and incubated at 37 °C and 5% CO_2_ for 30 minutes. Media was again removed and replenished with 30 mL phenol-red free Opti-MEM supplemented with 5 mM CT-TAMRA (Promega, G8251), except for no CT-TAMRA control wells, which were replenished with 30 mL phenol-red free Opti-MEM alone. Cells were then incubated at 37 °C and 5% CO_2_ for 30 minutes. Media was removed and cells were washed a final time with phenol-red free Opti-MEM, this time supplemented with 10% FBS + 1% Penicillin/Streptomycin and incubated at 37 °C and 5% CO_2_ for 30 minutes. Media was removed, and cells were washed once in 50 mL of 1X PBS and trypsinized with 12.5 mL of clear 0.25% Trypsin-EDTA (5X) (Gibco, 15400-054) per well. Cells were incubated at 37 °C and 5% CO_2_ for 15–20 minutes to ensure in complete dislodging of cells from the assay plate. Trypsin was then quenched with 12.5 mL of 50% FBS in 1X PBS. Flow cytometry was completed on an IntelliCyt iQue Screener PLUS equipped with ForeCyt acquisition software. Live, single cells were gated first for GFP expression, and GFP positive cells were then analyzed for mean fluorescence intensity of CT-TAMRA dye by double normalization to a no dye sample (0% red signal) and dye only sample (100% red signal). Data analysis was completed with FlowJo and GraphPad Prism 8 software.

### Surface Plasmon Resonance Experiments

SPR experiments were performed on a BioRad ProteOn XPR36 Interaction Array System. All compound stock solutions were diluted to desired final concentrations in SPR Buffer (20 mM Tris-HCl, pH 7.0, 150 mM NaCl, 0.005% Tween-20), and protein stock solutions were diluted into SPR Buffer supplemented with 1 mg/mL BSA. Biotinylated derivative of UNC7040, compound 39 (UNC7045) was made up as 150 nM stocks in SPR buffer, and immobilized at a flow rate of 30 μL/min and a contact time of 60 s onto a NeutrAvidin-containing ProteOn NLC sensor chip. Following a 30 min buffer blank in which SPR Buffer was switched to buffer supplemented with 1 mg/mL BSA, proteins (CBX2, CBX4, CBX6, CBX8, CDYL2) were flowed at a rate of 50 μL/min with a contact time of 200 s and a dissociation time of 800 s. Regeneration of the sensor chip in 0.1% SDS/5mM NaOH was completed between each protein sample at a flow rate of 30 μL/min for 120 s. Double referencing subtraction was done with buffer and protein blank channels to account for nonspecific binding to the sensor chip. Data were fit to a two-state binding model in which ka and kd parameters were fit as grouped, ka2, kd2, and RI parameters were fit locally, and all other parameters were fit globally.

### Protein Domain Microarray Analysis

The protein domain microarray was conducted by the Protein Array and Analysis Core (PAAC) at MD Anderson Cancer Center. A comprehensive library of human protein domains that potentially read methyllysine marks was cloned into a pGEX vector by Biomatik (Cambridge, Canada) using gene synthesis to best optimize the open reading frames for bacterial expression. Escherichia coli was used to express the protein domains as GST fusions, which were purified using glutathione-Sepharose beads. The recombinant domains were arrayed onto nitrocellulose-coated glass slides (Oncyte®Avid slides, Grace Bio-Labs, Bend, OR), using an Aushon 2470 pin microarrayer, as previously described (Espejo et al., 2002). Biotinylated compounds were pre-conjugated to streptavidin-Cy3 to generate fluorescent probes. Following probing, fluorescent signal was detected with a GenePix 4200A Microarray Scanner (Molecular Devices).

### Molecular Dynamics and Docking

Molecular dynamics (MD) simulations for all three systems (CBX8_CD_, CBX8_CD_:**21**, and CBX8_CD_:**22**) were performed using the Gromacs 2018.2 simulation package with CHARMM22 protein force field (Vanommeslaeghe et al., 2010). The 3D structures of CBX8_CD_ in complex with **21** and **22** were built by analogy from the crystal structure of CBX8 in complex with a chemically similar ligand UNC3866 (PDB: 5EQ0) (Stuckey et al., 2016) using the Maestro modeling suite (release 2016-2, Schrödinger, LLC: New York, NY). Both above structures served as starting points for MD simulations. CHARMM22 force field parameters for **21** and **22** were generated by Swissparam (Zoete et al., 2011). End caps were added to both termini of each protein. The protein complex was minimized in vacuum using steepest decent algorithm for 5,000 steps or until the maximum force of 1,000 kJ*mol^-1^*nm^-1^ was reached. The molecular systems were then solvated in TIP3P water (Mark and Nilsson, 2001), counterions were added to ensure the systems’ electric neutrality, and NaCl ions (0.15 M) were added by randomly replacing certain water molecules in order to mimic physiological conditions. An energy minimization with solvent was then performed, followed by a two-step equilibration: 5 ns in NVT ensemble at 310 K using the modified Berendsen thermostat and 5 ns in NPT ensemble at 1 atm (and 310 K) using the Parinello-Rahman pressure coupling (Nosé and Klein, 1983). All simulations were conducted using the Leapfrog integrator in periodic boundary conditions. The particle mesh Ewald algorithm (Essmann et al., 1995) controlled the long-range electrostatic interactions. Bonds involving hydrogen atoms were constrained using the linear constraint solver algorithm (LINCS) (Hess et al., 1997). The production simulations were performed in NVT ensemble. For each of the three systems, three independent ∼5 µs MD simulations were run. Molecular visualizations were produced using Maestro [Schrodinger, LLC]. MD trajectories were clustered and analyzed by means of the Pipeline Pilot data processing environment (v. 18.1.100.11, BIOVIA, 3dsbiovia.com). The input data (sets of the protein’s atomic coordinates and the backbone ϕ and ψ angles) were generated from the MD trajectories using custom Pipeline Pilot scripts (protein structures were centered and aligned using the Gromacs trjconv tool). The clustering technique used was *k-*means with Euclidian distance metrics. The cluster aggregation criteria were chosen so that root mean square distances (RMSD) between the cluster members would be on the order of 1 Å. Protein-DNA docking calculations were performed using the HADDOCK web service (van Zundert et al., 2016; Wassenaar et al., 2012). Twelve protein structures were selected for docking from the MD trajectories of the two simulated systems (CBX8_CD_:**21** and CBX8_CD_:**22**; 6 structures per systems). All selected structures were centroids of the most populated conformational clusters. The 3D structure of the DNA double helix (35 base pairs) was generated using the Discovery Studio 4.0 modeling suite (www.3dsbiovia.com). A set of default HADDOCK parameters was used for all docking simulations. The parameter file and all input and output HADDOCK files are available upon request.

### DLBCL cell proliferation assay

SUDHL4 cells were seeded at a cell density of 1x10^5 cells/mL in a 24 well plate (Corning® Costar®, 3526), with 0.5mL of cells per well. The vehicle control treated cells were dosed with 0.4% water or DMSO. Every 48 hours, the cells were counted on an automated Bio-Rad TC20™ cell counter with Trypan blue (Abcam, ab233465) and cell counting slides (1450015) to give the cell count (cells/mL) and cell viability (%). At each time point, the cells were split back to the original seeding density with fresh media, and the cells were re-dosed with compound or vehicle. The percent cell proliferation of cells treated with compound was calculated based on the total cell number expressed as split-adjusted viable cells, relative to the vehicle control at the same time point. The data is reported as the average of between 4 to 6 replicates ± standard deviation.

### CRC cell proliferation assays

HCT116, Caco2, LoVo and DLD-1 were seeded in 96-well Cell Carrier plates (Perkin Elmer) at 1500, 2000, 1200 and 1000 cells per well, respectively. The day after seeding (D0), cells were treated with various concentrations of UNC7263, UNC7040, UNC4976 or vehicle. Live and dead cell counts were measured daily over a 3-day period using the Operetta High Content Screening system (PerkinElmer). Prior to imaging, cells were stained with 5 μg/ml of Hoechst 33342 (nuclear dye) (Life Technologies) and 5 μg/ml of propidium iodine (vital dye) (Life Technologies) for 30 minutes. Cells were then segmented based upon the nuclear dye using the Harmony software (PerkinElmer). Propidium iodide intensity levels were calculated and cells were classified as ‘dead’ if their intensity was above the established threshold.

### CRC colony assay formation

To test the effect of CBX 8 inhibitors on the cell proliferation, we performed colony assays. For colony assay, HCT116 or LoVo cells were plated at the density of 500 cells per well in six well plates. After 24 hours, cells were treated with 10 or 20 mM UNC7040 or UNC7263. Depending on cell lines, after 1 to 2 weeks colonies were fixed with 100 % methanol for 20 minutes at room temperature, followed by rinsing with water. Next, colonies were stained with 0.5%crystal violet in 25 % methanol for 5 minutes at room temperature. Cells were washed with water to remove excess dye. Plates were inverted and left overnight for drying. Colonies were viewed using bright field microscopy. A non-overlapping cluster of minimum 50 cells were counted as a colony. All the treatments were done in triplicates.

### Chromatin Immunoprecipitation and quantitative PCR (ChIP-qPCR) and Next-Generation Sequencing (ChIP-seq)

25x10^6^ mES cells were collected, washed in once in 1x PBS and crosslinked with formaldehyde at a final concentration of 1 % for 7 min. The crosslinking was stopped on ice and with glycine at final 0.125 M concentration. The crosslinked cells were pelleted by centrifugation for 5 min at 1200g at 4 °C. Nuclei were prepared by washes with NP-Rinse buffer 1 (final: 10 mM Tris pH 8.0, 10 mM EDTA pH 8.0, 0.5 mM EGTA, 0.25% Triton X-100) followed by NP-Rinse buffer 2 (final: 10 mM Tris pH 8.0, 1 mM EDTA, 0.5 mM EGTA, 200 mM NaCl). Afterwards the cells were prepared for shearing by sonication by two washes with Covaris shearing buffer (final: 1 mM EDTA pH 8.0, 10 mM Tris-HCl pH 8.0, 0.1% SDS) and resuspension of the nuclei in 0.9 mL Covaris shearing buffer (with 1× protease inhibitors complete mini (Roche)). The nuclei were sonicated utilizing 15 ml Bioruptor tubes (Diagenode, C01020031) with 437.5 mg sonication beads (Diagenode, C03070001) for 6 cycles on a Bioruptor Pico sonicator. Lysates were incubated in 1x IP buffer (final: 50 mM HEPES/KOH pH 7.5, 300 mM NaCl, 1 mM EDTA, 1% Triton X-100, 0.1% DOC, 0.1% SDS), with following antibodies at 4 °C on a rotating wheel: H3K27me3 (Diagenode, C15410195), Ring1B (Cell Signaling, D22F2), Suz12 (Cell Signaling, D39F6), Cbx7 (Abcam, ab21873). ChIPs were washed 5x with 1x IP buffer (final: 50 mM HEPES/KOH pH 7.5, 300 mM NaCl, I mM EDTA, 1% Triton-X100, 0.1% DOC, 0.1% SDS), or 1.5x IP buffer for H3K27me3, followed by 3x with DOC buffer (10 mM Tris pH 8, 0.25 mM LiCl, 1 mM EDTA, 0.5% NP40, 0.5% DOC) and 1x with TE (+50 mM NaCl).

### qPCR Analysis

The PCIA extracted IP DNA was precipitated and quantified using a homemade EvaGreen based qPCR mix on a CFX Connect Real-Time PCR Detection System (BioRad). qPCR primers are listed in Table S4.

### ChIP-seq Library Preparation

Libraries were prepared using the NEXTflex ChIP-Seq kit (Bio Scientific) following the “No size-selection cleanup” protocol. Each sample of ChIPed DNA was end-repaired and ligated to unique barcoded adaptors to produce individual libraries. Libraries corresponding to samples to be directly compared to each other (e.g. DMSO vs UNC7040) were pooled together and purified using 1 volume of Agencourt AMPure XP (Beckman Coulter). The pooled libraries were eluted with 23 µL of elution buffer (NEXTflex ChIP-Seq kit) and amplified using the KAPA Real-Time Library Amplification Kit (KAPABiosystems) following the kit instructions. Finally, the amplified libraries were size-selected and purified using 0.9x volume of Agencourt AMPure XP (Beckman Coulter). Library quality control including determination of average size and concentration was performed prior to sequencing by commercial Next Generation Sequencing providers. NGS libaries were eventually sequenced as 150 bp paired-end reads on the Illumina HiSeq platform.

### ChIP-seq Data Analysis

#### Processing and mapping of raw reads

The raw reads of ChIP-seq were mapped to the custom concatenated human (hg38) and spike-in mouse (mm10) genome sequences using bowtie 2 with “–no-mixed” and “no-discordant” options (Langmead and Salzberg, 2012). Subsequently, low quality reads were removed using SAMtools (Li et al., 2009), duplicated reads were discarded with the Picard toolkit (http://broadinstitute.github.io/picard/) and only unique mapped reads were retained for subsequent analysis.

#### Data visualization

For visualization uniquely mapped human reads were normalized by random subsampling using calibration factors calculated from the corresponding mm10 spike-in reads as described previously (Fursova et al., 2019) using samtools . High correlation between replicates was confirmed using multiBamSummary and plotCorrelation functions from deepTools (Ramírez et al., 2016) before merging for visualization and downstream analysis. Genome coverage tracks (bigWig files) were generated with MACS2’ pileup function (Zhang et al., 2008) and heatmaps and profile plots were generated with deepTools (Ramírez et al., 2016).

#### Peak calling

Peaks were called on each replicate independently using MACS2 (Zhang et al., 2008) with -broad option and q-value cutoff of 0.1. Only peaks called in both replicates were retained for downstream analysis.

### RNA-seq Library Preparation

For RNA-seq sample preparation, 5-10x10^6 SUDHL-4 or LoVo cells (untreated, UNC7263 or UNC7040 treated in triplicates) were trypsinized and collected by centrifugation for 5 min at 500 g. Subsequently, cell pellets were washed 1× with PBS and collected in 1X DNA/RNA protection reagent (Monarch Total RNA Miniprep Kit, NEB). Cells were lysed and total RNA was isolated following the mammalian cell protocol inlcuding on-column DNase I treatment. Total RNA samples were submitted to Novogene Co. for quality control and library preparation, applying poly-A enrichment and using the NEBNext Ultra II RNA Library Prep kit (NEB), indexed using NEBNext Multiplex Oligos (NEB). Final libraries were sequenced as 150 bp paired-end reads on the Illumina HiSeq platform.

### RNA-seq Data Analysis

Raw paired RNA-seq reads were aligned to the hg38 genome sequences using STAR-2.6.1c (Dobin et al., 2013). Overlap of aligned reads in bam format with genes was performed using HTSeq count function (Anders et al., 2015) with stranded=no option and the GRCh38 version 101 GTF file. The HTseq count matrix was pre-filtered to remove any genes with less than 10 reads. Differential gene expression analysis was performed using DESeq2 (Love et al., 2014) using “apeglm” method (Zhu et al., 2019) for LFC shrinkage. We applied a threshold of p-adj <= 0.05 and fold change >= 0.5 or <=-0.5 to consider gene expression changes significant. Log2 fold change values were visualized in volcano plots using R, ggplot2 and GraphPad prism. To assign intergenic cis-regulatory regulatory elements identified by ChIP-seq with associated genes we utilized the Genomic Regions Enrichment of Annotations Tool (GREAT) (McLean et al., 2010).

### Co-immunoprecipitation

Whole cell protein extract from CBX8 reporter mESCs (45 × 10^6) was obtained by lysis in 500 µl lysis buffer (final: 20 mM Tris-HCl pH 7.5, 150 mM NaCl, 2 mM MgCl_2_, 10% Glycerol, 1 mM DTT, 1 mM PMSF, 0.2% NP-40, 1× Complete Mini protease inhibitor). Lysate was homogenized by sonication in a Diagenode Bioruptor Pico for 20 cycles followed by rotation for 3 hours at 4 °C. After 30 min centrifugation at 4 °C protein concentration of the lysate was determined by Bradford assay (Biorad). In parallel, 30 µl Protein G coupled Dynabeads (Thermo Fisher Scientific) were prepared for each IP reaction as follows: 3x wash in lysis buffer, incubation with 1.5 µg FLAG antibody (Sigma Aldrich, F1804) for 3 hours at 4 °C, 1× wash in lysis buffer and finally resuspension in 30 µl of lysis buffer. For each IP, 30 µl of pre-bound Dynabeads were incubated with 1 mg protein extract in a final volume of 500 µl overnight at 4 °C. Finally, beads were washed four times with lysis buffer and proteins were eluted at 95 °C in SDS sample buffer and analyzed by Western blot.

Co-immunoprecipitation samples and corresponding 2% Input samples were run on Novex Life Technology NuPAGE 4–12% Bis-Tris gels in Invitrogen NuPAGE MES SDS Running Buffer and transferred on a Merck Chemicals Immobilon-FL Membrane (PVDF 0.45 µm). The membrane was blocked (5% non-fat dry milk in 1× PBS, 0.1% Tween 20) and incubated in 5% non-fat dry milk in 1× PBS and 0.1% Tween 20 with the primary antibodies (PHC1 Cell Signaling #13505 1:1000; Rybp Cell Signaling D8J7W 1:1000; SUZ12 Cell Signaling (D39F6) 1:1000; Cbx7 Merck Millipore 07-981 1:1000; Cbx8 Bethyl Laboratories A300-882A 1:1000; Ring1B Cell Signaling D22F2 1:1000). Finally, the membrane was incubated with corresponding secondary IRDye 800CW Goat anti-Rabbit IgG (H+L) (LICOR) or IRDye 680RD Goat anti-Mouse IgG (H+L) (LICOR) and imaged by an Odyssey CLx Near-Infrared Imaging System (LICOR).

---

**Figure S1.**
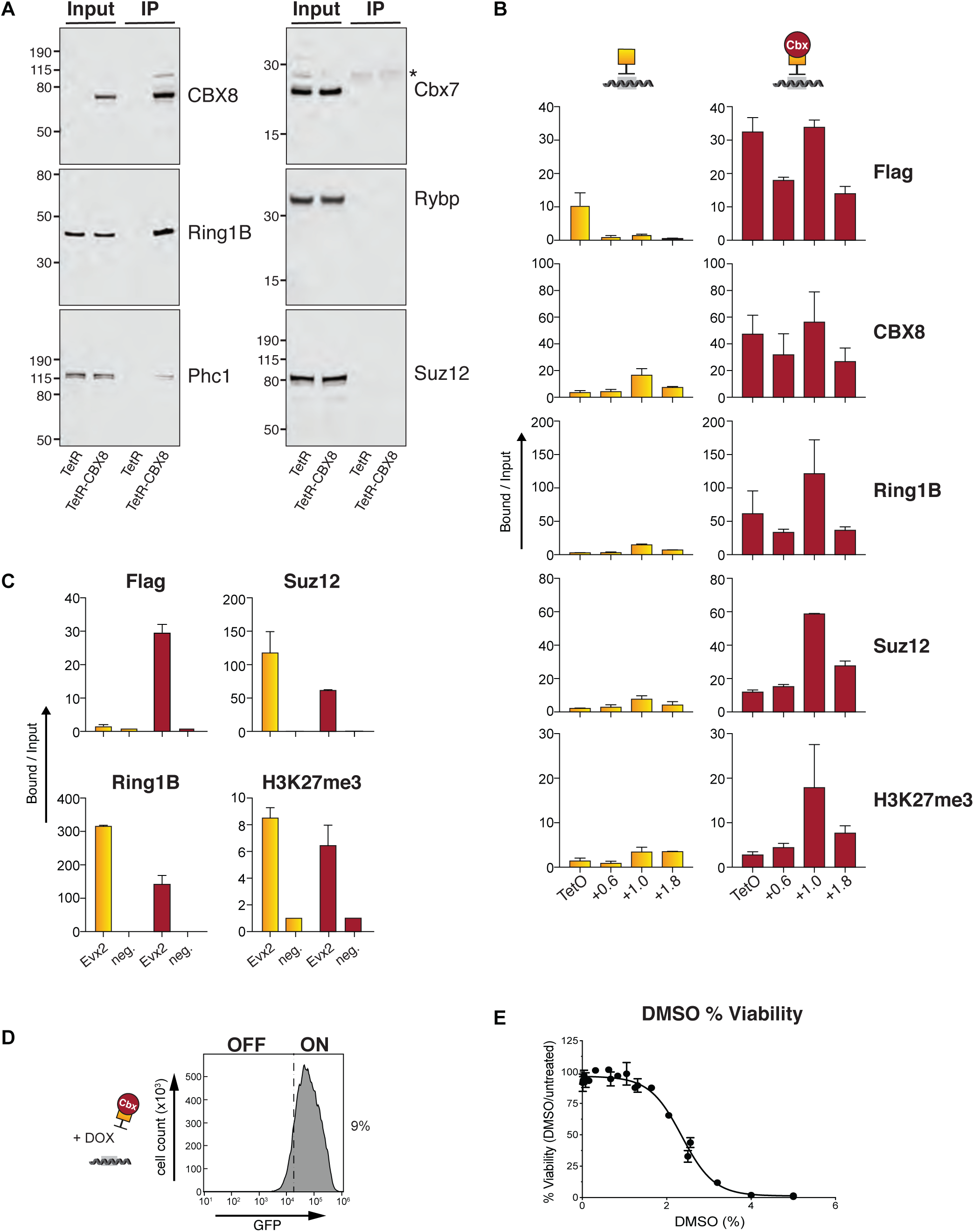
TetR-CBX8 associates with endogenous cPRC1 components and triggers formation of a repressive Polycomb chromatin domain at the reporter locus in mESCs. (A) Co-immunoprecipitation analysis of FLAG-epitope-tagged TetR-CBX8 in reporter mESCs reveals interaction with cPRC1 subunits Ring1B and Phc1 but not vPRC1-specific Rybp or PRC2 subunit Suz12. TetR-CBX8 association with RING1B excludes endogenous Cbx7. ChIP-qPCR analysis shows relative enrichments of FLAG-TetR alone or fused to CBX8, PcG proteins and H3K27me3 at and downstream of the TetO DNA binding sites (TetO) (B) and at the Evx2 promoter (positive control) and at IAP (negative control) (C). ChIP enrichments are normalized to IAP. Data are mean ± SD (error bars) of at least two independent experiments. (D) Flow cytometry histogram shows GFP expression in TetR-CBX8 expressing reporter mESCs after DOX treatment for four days. Percentage indicates fraction of GFP-negative reporter mESCs. (E) Viability of reporter mESCs in response to DMSO treatment.

**Figure S2.**
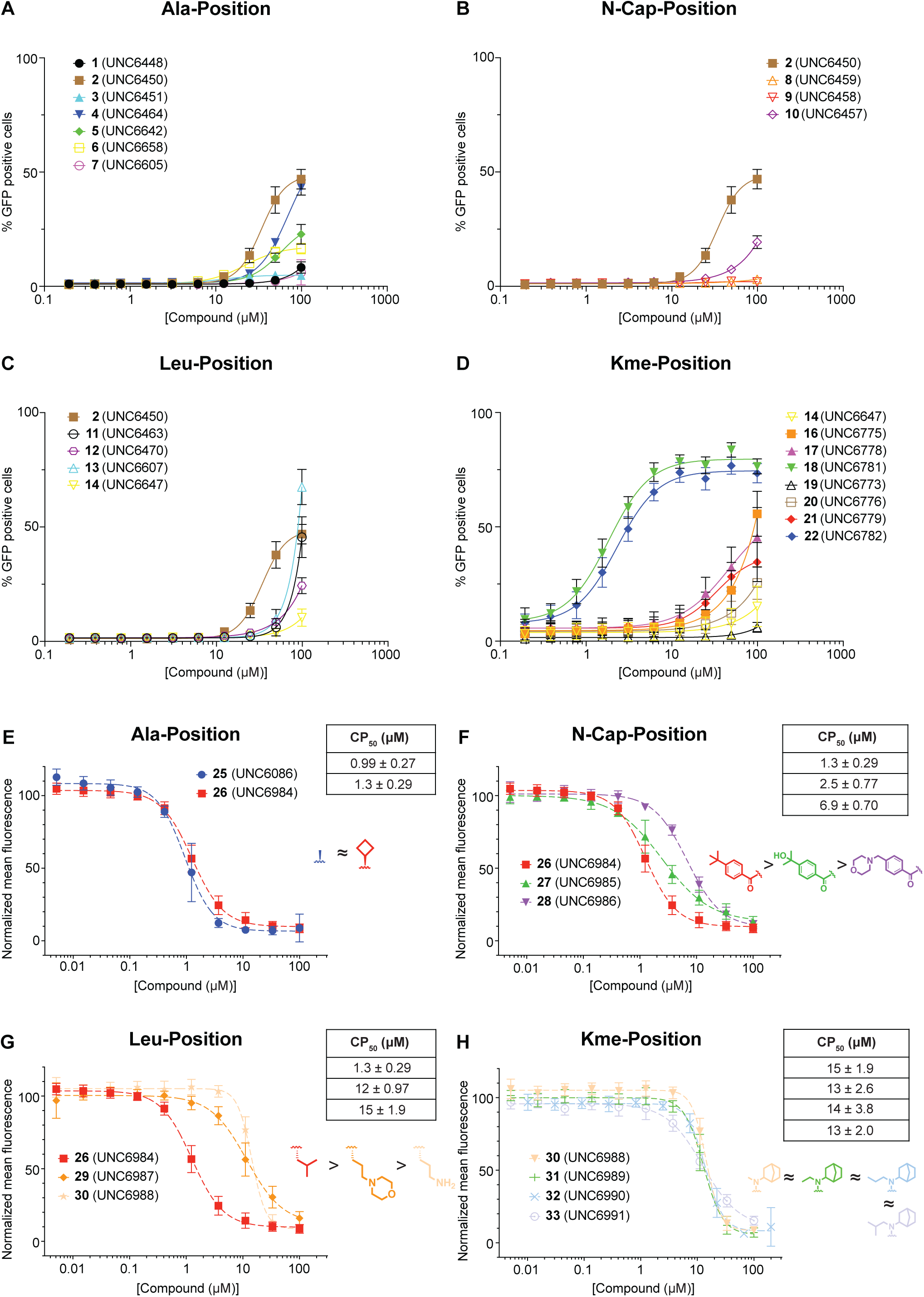
Cellular activity screening data and cell permeability assay results of each amino acid modification. CBX8 GFP reporter assay results of modified compounds at (A) Ala- position, (B) N-cap-position, (C) Leu-position, and (D) Lysine mimetics-position. CAPA assay results of modified compounds at (E) Ala-position, (F) N-cap-position, (G) Leu-position, and (H) Lysine mimetics-position. Data shown are mean ± SD, n = 9.

**Figure S3.**
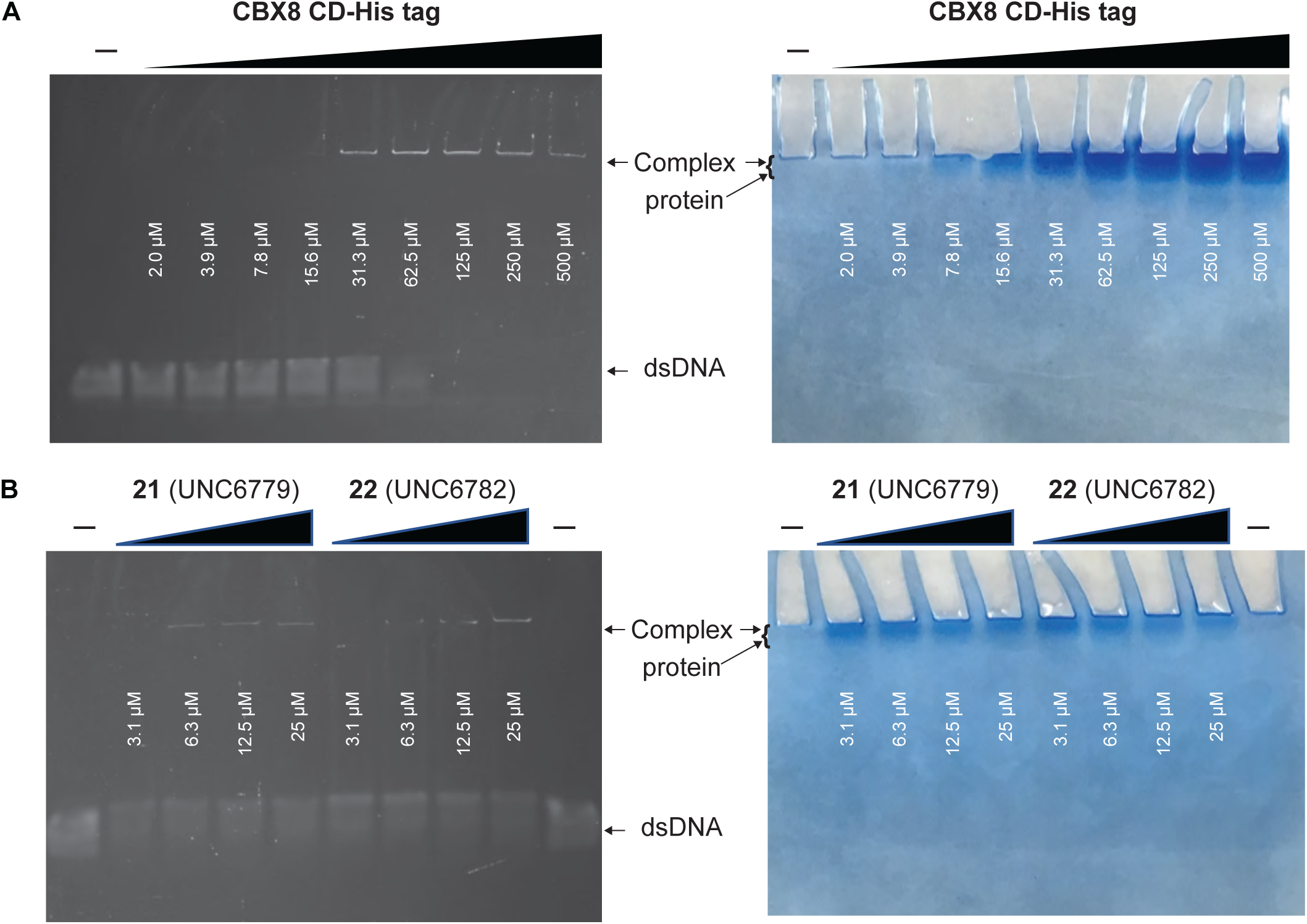
Comparison of electrophoretic mobility shift assay (EMSA) results between compound 21 and 22. (A) Fluorescent image of EMSA protein titration of CBX8 chromodomain (left) and the Coomassie staining of the gel (right), (B) Fluorescent image of EMSA compound titration of 21 and 22 (left) and the Coomassie staining of the gel (right).

**Figure S4.**
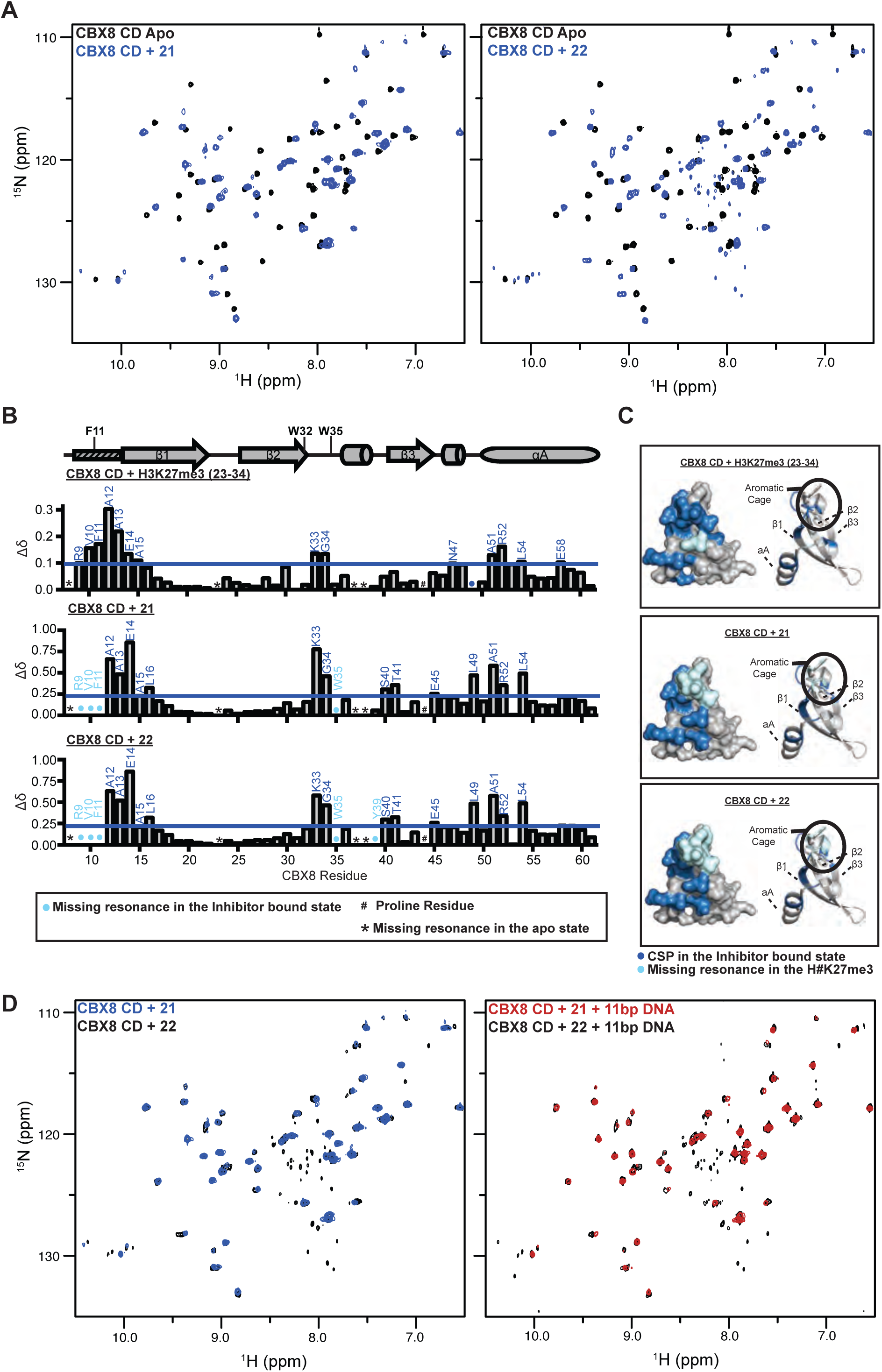
CBX8-CD residues with CSPs upon binding to H3K27me3, 21, and 22. (A) Full ^1^H-^15^N-HSQC overlays for ^15^N-CBX8-CD upon addition of compound 21 (left) or 22 (right). (B) Normalized CSP (Δδ) between H3K27me3-bound (top), 21-bound (middle), and 22-bound (bottom) spectra are plotted against CBX8 residue number. (C) Residues with significant CSPs upon addition of H3K27me3 peptide, compound 21, and compound 22 plotted onto a cartoon and surface representation of the CD and colored blue. Missing resonance is colored light blue.

**Figure S5.**
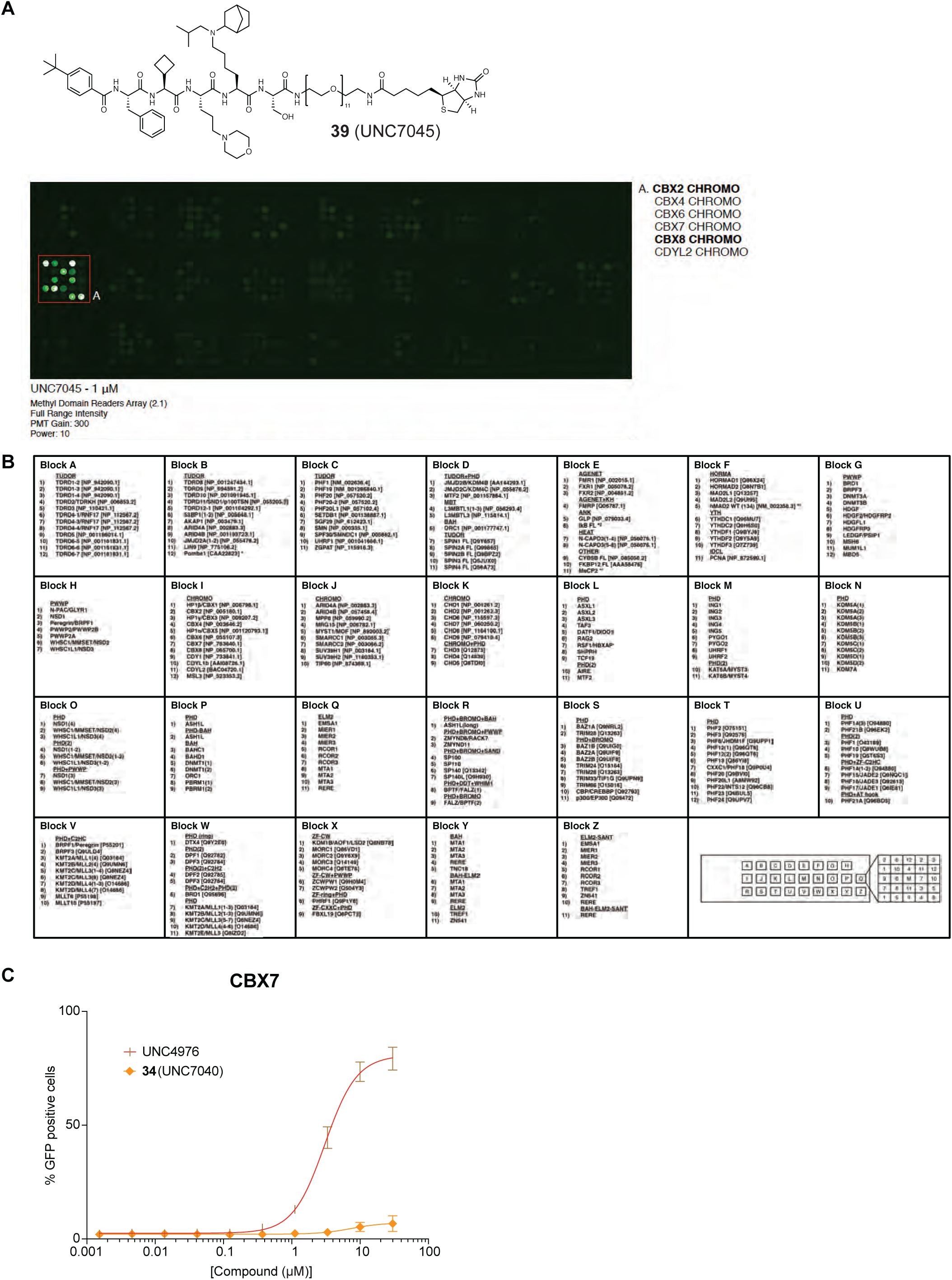
*In vitro* and cellular selectivity profiling of compound 34. (A) Microarray result of selectivity profiling of **39** (biotinylated **34**). (B) Complete list of histone Methyl-lysine reader domains evaluated on the protein microarray screened. Mapping of individual reader domains is described by the legend in the bottom right corner. (C) Cell reporter selectivity result of **34** using the CBX7 GFP reporter assay. n = 9.

**Figure S6.**
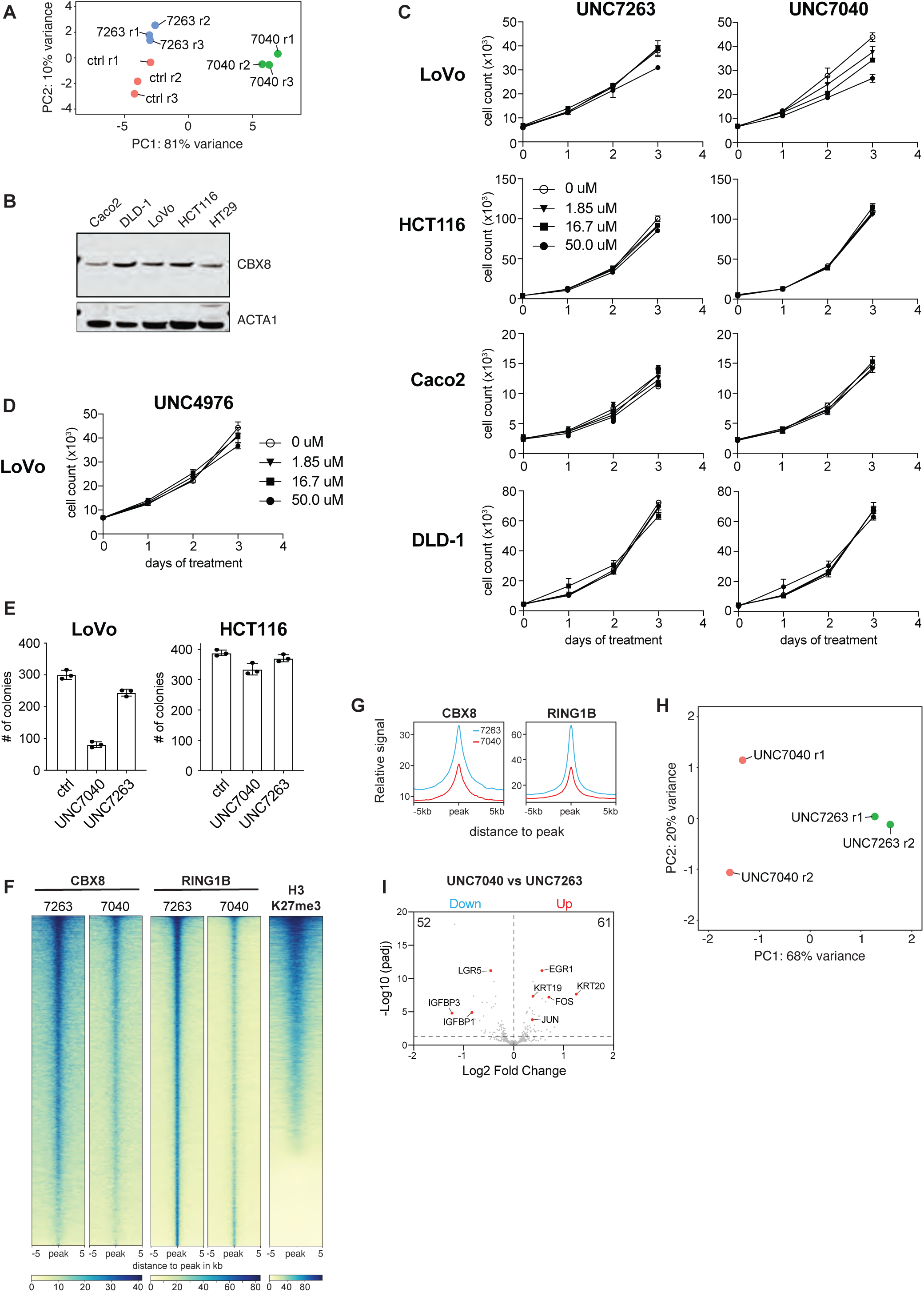
UNC7040 causes reduced colorectal cancer cell proliferation and activation of differentiation genes by blocking canonical PRC1 targeting via CBX8. (A) Principle component analysis (PCA) of three independent replicates of RNAseq in SUDHL4 cells treated with DMSO, UNC7040 or UNC7263. (B) Immuno blot analysis of CBX8 expression in panel of CRC cell lines. ACTA1 serves as loading control. (C) Effects of treatment with DMSO, UNC7263 or UNC7040 on proliferation of different CRC cell lines. (E) Effects of treatment with DMSO, UNC7263 or UNC7040 on colony formation in LoVo and HCT116 cells. (F) Heatmaps of CBX8 and RING1B ChIPseq enrichment in LoVo cells treated with UNC7263 at 20 μM or UNC7040 at 20 μM. Also shown is H3K27me3 in UNC7263-treated LoVo cells. The signals from two independent cChIPseq experiments are centered around RING1B peaks ± 5 kb and plotted as spike-in normalized mapped read counts. (G) Meta plots display CBX8 and RING1B occupancy in LoVo cells in response to treatment with UNC7263 (blue) or UNC7040 (red). (I) Volcano plots show gene expression changes in LoVo cells treated with UNC7040 relative to UNC7263. Numbers in represent repressed (blue) and upregulated (red) genes. Differential gene expression (P_adj_ < 0.05; LFC <-0.5 and >0.5, respectively) was calculated from three independent replicates. Highlighted are genes associated with CRC stem cell proliferation and differentiation. (H) PCA of two independent replicates of RNA-seq in LoVo cells treated with UNC7040 or UNC7263.

**Table S1.**
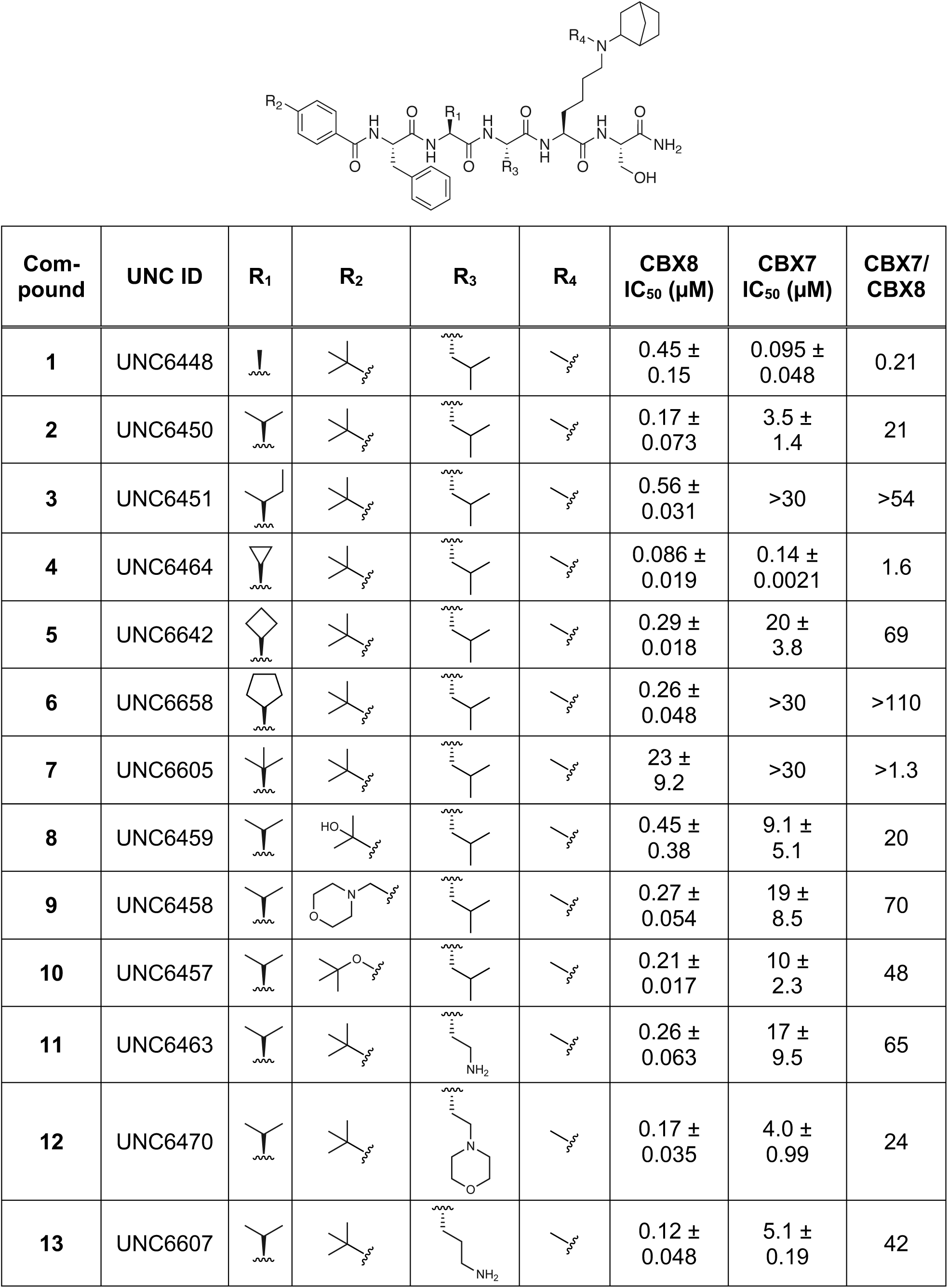
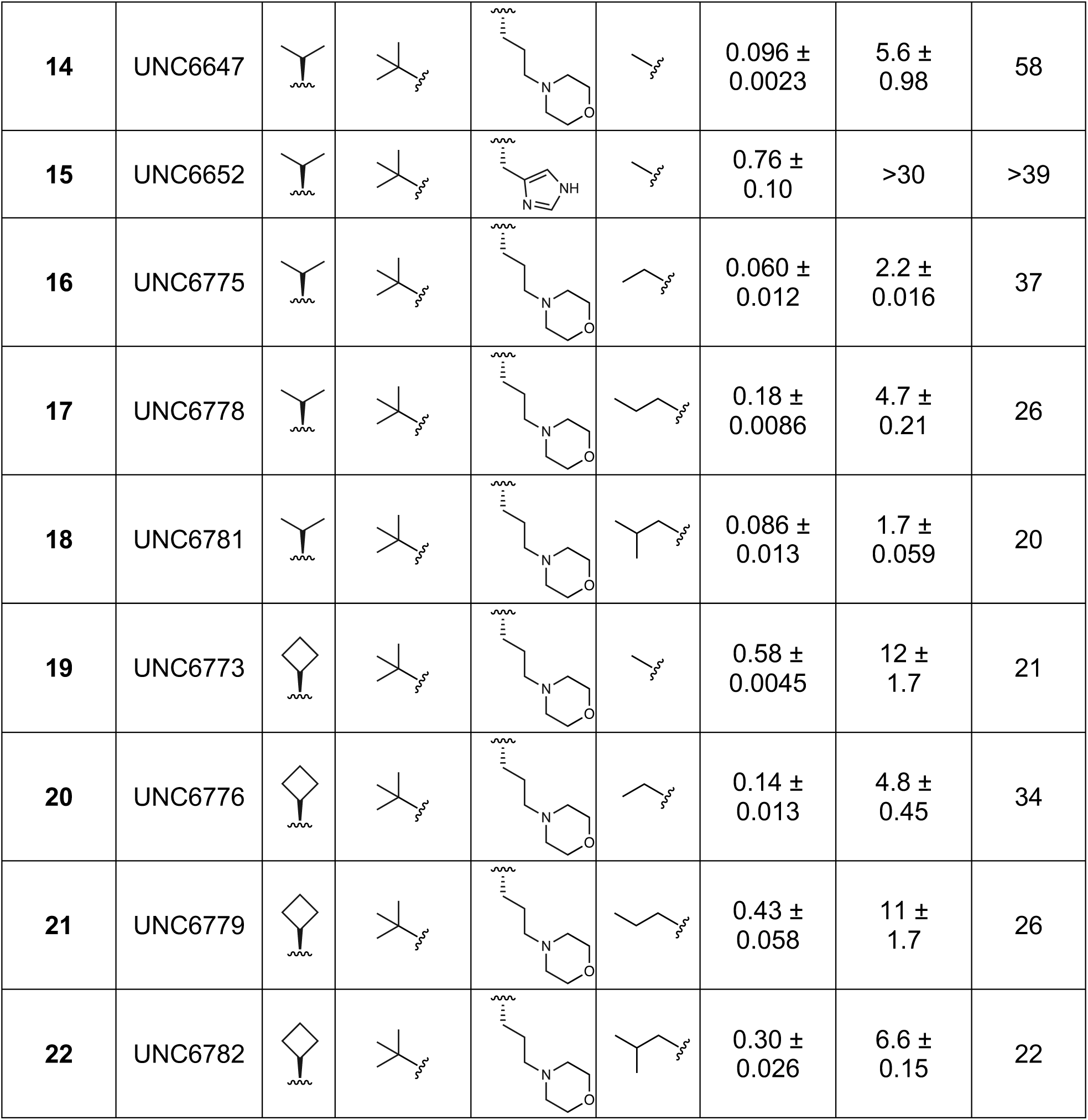
Competitive displacement IC_50_’s in TR-FRET assay of designed CBX8 antagonists for CBX7 and CBX8 and their improved CBX8 selectivity over CBX7. Data shown are mean ± SD, n = 6.

| Com-<br>pound | UNC ID | R<sub>1</sub> | R<sub>2</sub> | R<sub>3</sub> | R<sub>4</sub> | CBX8<br>IC<sub>50</sub> (μM) | CBX7<br>IC<sub>50</sub> (μM) | CBX7/<br>CBX8 |
| --- | --- | --- | --- | --- | --- | --- | --- | --- |
| 1 | UNC6448 |  |  |  |  | 0.45 ± 0.15 | 0.095 ± 0.048 | 0.21 |
| 2 | UNC6450 |  |  |  |  | 0.17 ± 0.073 | 3.5 ± 1.4 | 21 |
| 3 | UNC6451 |  |  |  |  | 0.56 ± 0.031 | >30 | >54 |
| 4 | UNC6464 |  |  |  |  | 0.086 ± 0.019 | 0.14 ± 0.0021 | 1.6 |
| 5 | UNC6642 |  |  |  |  | 0.29 ± 0.018 | 20 ± 3.8 | 69 |
| 6 | UNC6658 |  |  |  |  | 0.26 ± 0.048 | >30 | >110 |
| 7 | UNC6605 |  |  |  |  | 23 ± 9.2 | >30 | >1.3 |
| 8 | UNC6459 |  |  |  |  | 0.45 ± 0.38 | 9.1 ± 5.1 | 20 |
| 9 | UNC6458 |  |  |  |  | 0.27 ± 0.054 | 19 ± 8.5 | 70 |
| 10 | UNC6457 |  |  |  |  | 0.21 ± 0.017 | 10 ± 2.3 | 48 |
| 11 | UNC6463 |  |  |  |  | 0.26 ± 0.063 | 17 ± 9.5 | 65 |
| 12 | UNC6470 |  |  |  |  | 0.17 ± 0.035 | 4.0 ± 0.99 | 24 |
| 13 | UNC6607 |  |  |  |  | 0.12 ± 0.048 | 5.1 ± 0.19 | 42 |
| 14 | UNC6647 | | | | | $0.096 \pm 0.0023$ | $5.6 \pm 0.98$ | 58 |
| 15 | UNC6652 | | | | | $0.76 \pm 0.10$ | >30 | >39 |
| 16 | UNC6775 | | | | | $0.060 \pm 0.012$ | $2.2 \pm 0.016$ | 37 |
| 17 | UNC6778 | | | | | $0.18 \pm 0.0086$ | $4.7 \pm 0.21$ | 26 |
| 18 | UNC6781 | | | | | $0.086 \pm 0.013$ | $1.7 \pm 0.059$ | 20 |
| 19 | UNC6773 | | | | | $0.58 \pm 0.0045$ | $12 \pm 1.7$ | 21 |
| 20 | UNC6776 | | | | | $0.14 \pm 0.013$ | $4.8 \pm 0.45$ | 34 |
| 21 | UNC6779 | | | | | $0.43 \pm 0.058$ | $11 \pm 1.7$ | 26 |
| 22 | UNC6782 | | | | | $0.30 \pm 0.026$ | $6.6 \pm 0.15$ | 22 |

**Table S2.** Halo-tagged compounds for cell permeability assay.

| Compound | UNC ID | R<sub>1</sub> | R<sub>2</sub> | R<sub>3</sub> | R<sub>4</sub> |
| --- | --- | --- | --- | --- | --- |
| **25** | UNC6086 | *t*-Bu | methyl | isobutyl | methyl |
| **26** | UNC6984 | *t*-Bu | cyclobutyl | isobutyl | methyl |
| **27** | UNC6985 | 2-hydroxypropyl | cyclobutyl | isobutyl | methyl |
| **28** | UNC6986 | morpholinomethyl | cyclobutyl | isobutyl | methyl |
| **29** | UNC6987 | *t*-Bu | cyclobutyl | 4-morpholinopropyl | methyl |
| **30** | UNC6988 | *t*-Bu | cyclobutyl | aminopropyl | methyl |
| **31** | UNC6989 | *t*-Bu | cyclobutyl | aminopropyl | ethyl |
| **32** | UNC6990 | *t*-Bu | cyclobutyl | aminopropyl | propyl |
| **33** | UNC6991 | *t*-Bu | cyclobutyl | aminopropyl | isobutyl |

